# Innervation and cargo-specific axonal transport impairments in FUS-ALS mice with gain and loss of function

**DOI:** 10.1101/2025.06.16.659513

**Authors:** Rebecca L Simkin, Andrew P Tosolini, Alla Mikheenko, Agnieszka M Ule, Wenanlan Jin, Bernadett Kalmar, Mhoriam Ahmed, Qiuhan Lang, Franciane Lirot, Martha F McLaughlin, Georgia Price, David Villarroel-Campos, Linda Greensmith, James N Sleigh, Thomas J Cunningham, Giampietro Schiavo, Elizabeth M C Fisher, Pietro Fratta, Nicol Birsa

## Abstract

Mutations in the RNA-binding protein FUS lead to nuclear depletion and cytoplasmic mislocalisation of the protein and cause amyotrophic lateral sclerosis (ALS). Using a novel FUS-ALS mouse model, we found that adult mutant mice develop loss of function transcriptomic alterations, along with aberrant cytoplasmic partitioning associated with translatome deficits. Neuromuscular junction innervation was selectively impaired in FUS-ALS females; however, reinnervation following sciatic nerve crush was equally perturbed in both sexes. Additionally, we observed cargo-specific axonal transport alterations, a process critical for neuronal maintenance. *In vivo* mitochondrial transport was impaired across FUS-ALS mice, whereas the transport of signalling endosomes was selectively disrupted in mutant females. Altogether, our findings identify broader and sex-dependent motor neuron dysfunction in FUS-ALS, emphasise the link between endosomal transport impairments and denervation in disease, and establish FUS-ALS mice as a valuable model for investigating early cellular impairments driving ALS pathology.

## Introduction

Amyotrophic lateral sclerosis (ALS) is a fatal neurodegenerative disease characterised by the loss of upper and lower motor neurons, resulting in the progressive degeneration of the neuromuscular system and ultimately death from respiratory failure. Mutations in the RNA binding protein (RBP) FUS cause ∼ 5% of familial ALS and are often associated with aggressive, early onset cases of the disease ^1^. These mutations lead to nuclear depletion and cytoplasmic mislocalisation of the protein, resulting in the formation of pathological inclusions that are a hallmark of FUS-ALS. In addition to familial FUS-ALS cases, FUS mislocalisation has been identified in sporadic ALS ^2^, and FUS inclusions are a hallmark of the third most common form of frontotemporal dementia (FUS-frontotemporal lobar degeneration, FTLD; ∼ 5-10% of cases), albeit in the absence of FUS disease-causative mutations^3–6^.

FUS nuclear loss of function leads to widespread transcriptional changes that occur in combination with cytoplasmic toxic gain of function mechanisms, including aberrant phase-separation, translational deficits and altered stress response ^7–9^. Cultured mouse and human motor neurons, along with the use of homozygous lines to accentuate subtle pathological features, have been pivotal for identifying other pathological phenotypes caused by FUS-ALS mutants, such as impaired phase-separation, as well as defects in axonal transport and axonal outgrowth ^10–16^. Despite these advances, *in vivo* investigations in adult mice, which are critical for understanding the mechanisms underlying the pathogenesis of a late onset disease like ALS, have been hindered by several issues. For example, the tight dose-dependent autoregulation of FUS, the confounding studies in overexpressing transgenics, the perinatal lethality of FUS-ALS homozygous mice and the slowly progressive phenotype of heterozygous knock-in mice ^17–19^ have prevented the investigation of subtle, dose-dependent phenotypes within a physiologically-relevant context.

To overcome these challenges, we generated a novel homozygous mouse model that allowed us to investigate the effect of FUSALS mutations *in vivo*. Δ14FUS was identified in a case of aggressive, early onset ALS, and is caused by a mutation in the splice acceptor site of intron 13, which leads to skipping of exon 14 and a frameshift in exon 15 ^20^. This mutation, along with the humanisation of exon 15, recapitulates the frameshifted C-terminus found in the human mutant protein and is ubiquitously expressed at physiological levels ^17^. Δ14FUS leads to the complete ablation of the FUS nuclear localisation signal (NLS), resulting in the mislocalisation of the mutant protein in heterozygous animals ^10,17^. Similar to FUS knockout mice, Δ14FUS homozygotes on a C57BL/6J background are perinatally lethal; however, we found that C57BL/6J × DBA/2J intercrossing produces viable F1 progeny homozygous for the Δ14FUS mutation ^21^.

This new Δ14FUS homozygous knock-in model enabled us to dissect, in adult mice, the impact of FUS-ALS loss and gain of function on RNA metabolism, including expression, splicing and translation, and the effect of Δ14FUS expression on the neuromuscular system. At cellular level, we showed the aberrant, FUS-induced phase-separation of FMRP in spinal motor neurons, and the reduced translation of FMRP-bound RNAs that we previously identified in embryonic models ^10^. We found that NMJ innervation defects and *in vivo* signalling endosome transport disruption were selectively impaired in female, but not male, *Fus*^*Δ14/Δ14*^ mice, while mitochondrial transport and reinnervation following axonal damage were defective in a FUS-dependent manner for both sexes. Overall, our data strengthens the coupling between signalling endosome transport deficits and denervation in ALS, while highlighting broader transport deficits that may impact disease onset and pathogenesis.

## Results

### 1. Δ14FUS expression results in transcriptional alterations in adult *Fus*^*Δ14/Δ14*^ mice, compatible with nuclear loss of function

*Fus*^*Δ14/Δ14*^ adult mice were obtained as F1 progeny from *Fus*^*+/Δ14*^ C57BL/6J × *Fus*^*+/Δ14*^ DBA/2J intercrossing, as previously described ^21^. Due to the reported occurrence of fatal seizures in homozygous females from 3 months of age ^21^, experiments in this study were conducted by this endpoint. Δ14FUS was mislocalised in spinal motor neurons from *Fus*^*Δ14/Δ14*^ mice, as shown by the decreased nuclear/cytoplasmic ratio compared to *Fus*^*+/+*^ neurons (**Fig. 1A,B**; nuclear/cytoplasmic ratio in *Fus*^*+/+*^ = 2.72 ± 0.36, *Fus*^*+/Δ14*^ = 1.58 ± 0.16, *Fus*^*Δ14/Δ14*^ = 0.69 ± 0.11, *n* = 4). Additionally, mutant FUS localises to peripheral axons *in vivo*, showing a dosedependent enrichment in sciatic nerve axons of *Fus*^*+/Δ14*^ and *Fus*^*Δ14/Δ14*^ mutants compared to *Fus*^*+/+*^ controls (**Fig. 1C,D**; axonal *Fus*^*+/+*^= 2.37 ± 0.24, *Fus*^*+/Δ14*^ = 4.69 ± 0.48, *Fus*^*Δ14/Δ14*^= 9.33 ± 0.76, *n* = 6).

**Fig. 1.**
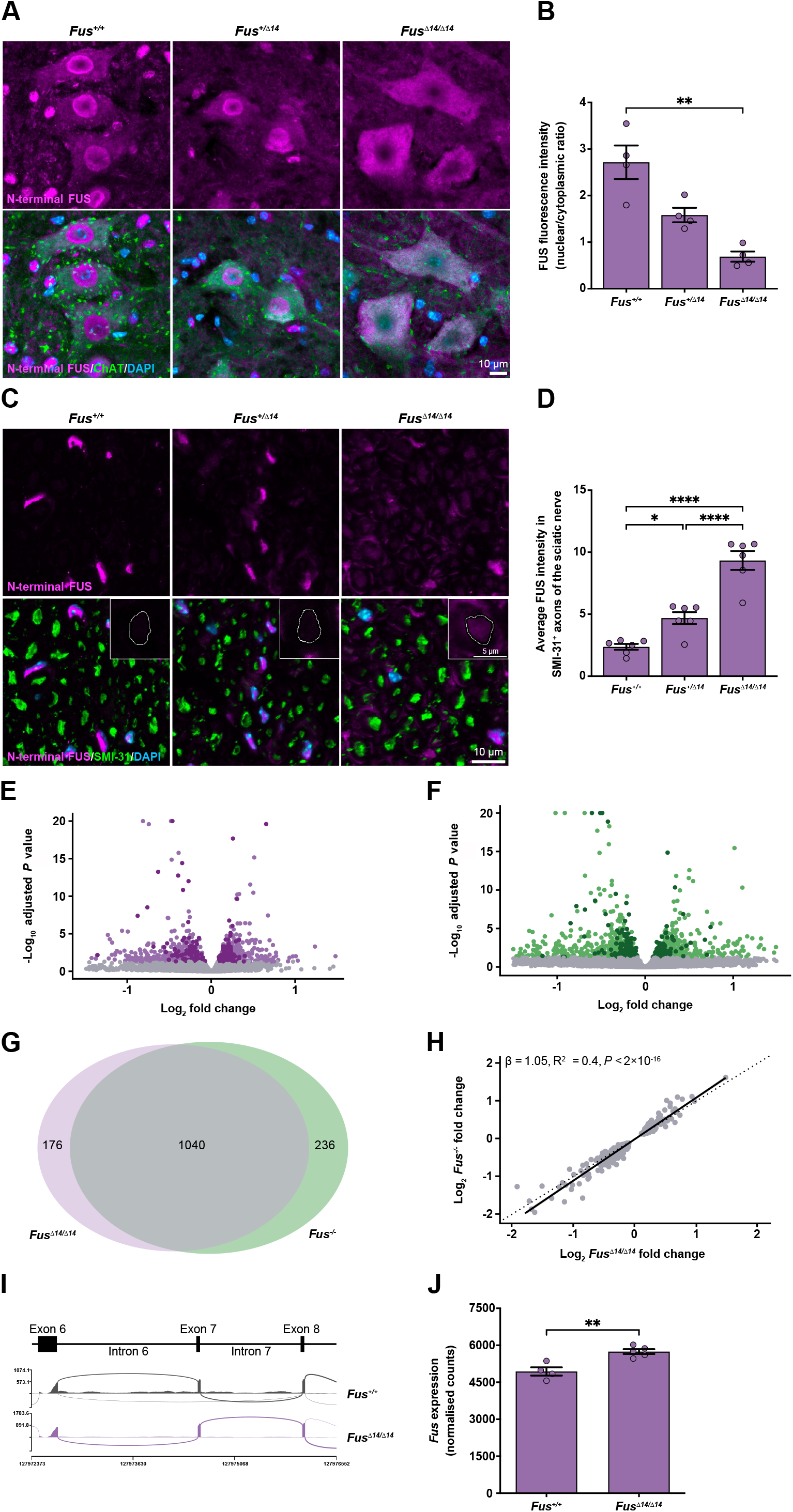
Δ14FUS is mislocalised in a dose-dependent manner in adult mice, resulting in nuclear loss of function. (**A**) Representative maximum z-stack projection images of *Fus*^*+/+*^ (left), *Fus*^*+/Δ14*^ (centre) and *Fus*^*Δ14/Δ14*^ (right) motor neurons in lumbar (L)1-L6 spinal cord. FUS is labelled using an N-terminal antibody recognising both wild-type and Δ14 mutant forms (magenta); cholinergic neurons are marked with ChAT (green), and nuclei with DAPI (blue). Scale bar = 10 µm. (**B**) Δ14FUS is mislocalised in *Fus*^*+/Δ14*^ and *Fus*^*Δ14/Δ14*^ motor neurons, as presented by the decreased ratio between nuclear and cytoplasmic FUS average intensities (P = 0.0005, Kruskal-Wallis). \*\**P* < 0.01, Dunn’s multiple comparisons. *n* = 4. (**C**) Representative maximum z-stack projection images of *Fus*^*+/+*^ (left), *Fus*^*+/Δ14*^ (centre) and *Fus*^*Δ14/Δ14*^ (right) sciatic nerve. Transverse sections were labelled with antibodies against phospho-neurofilament H (SMI-31; green) and N-terminal FUS (magenta), with nuclei counterstained using DAPI (blue). Inserts show examples of FUS labelling in single axon sections. Scale bars = 10 µm for larger images; 5 µm for inserts. (**D**) Image quantification reveals a dose-dependent increase in FUS levels within SMI-31 positive axons of *Fus*^*+/Δ14*^ and *Fus*^*Δ14/Δ14*^ sciatic nerve (P < 0.0001, one-way ANOVA). *P < 0.05, ****P < 0.0001, Bonferroni’s multiple comparisons. *n* = 6. (**E**) Volcano plot showing differentially expressed genes in *Fus*^*Δ14/Δ14*^ versus *Fus*^*+/+*^ datasets at 3 months. Known FUS targets are presented as darker dots, and non-significant changes (adj. *P* ≥ 0.05) are depicted in grey. (**F**) Volcano plot showing differentially expressed genes in *Fus*^*-/-*^ versus *Fus*^*+/+*^ datasets at 3 months. Known FUS targets are presented as darker dots, and nonsignificant changes (adj. P ≥ 0.05) are depicted in grey. (**G**) Venn diagrams illustrating the overlap (grey) between differentially expressed genes identified in *Fus*^*Δ14/Δ14*^ versus *Fus*^*+/+*^ (purple) and *Fus*^*-/-*^ versus *Fus*^*+/+*^ datasets (green). (**H**) Scatter plot showing the correlation between log^2^ fold change values of differentially expressed genes (adj. *P* < 0.05) in *Fus*^*Δ14/Δ14*^ versus *Fus*^*+/+*^ (x-axis) and *Fus*^*-/-*^ versus *Fus*^*+/+*^ (yaxis) datasets. (**I**) Schematic and sashimi plot showing decreased intron 6 and 7 reads in *Fus*^*Δ14/Δ14*^ versus *Fus*^*+/+*^ spinal cord samples. (**J**) *Fus* RNA counts reveal that expression of the mature transcript is significantly higher in *Fus*^*Δ14/Δ14*^ versus *Fus*^*+/+*^ datasets (P = 0.0032, unpaired t-test). *n* = 4-5.

To assess the transcriptional alterations resulting from loss of nuclear FUS, we performed high-depth RNA sequencing (RNA-seq) on spinal cord samples from 1 and 3 month-old male *Fus*^*+/+*^ and *Fus*^*Δ14/Δ14*^ mice. Differential gene expression analysis revealed widespread changes, with 1,200 differentially expressed genes (DEGs) in 1 month and 1,216 DEGs in 3 month *Fus*^*Δ14/Δ14*^ mutants compared to sex- and aged-matched *Fus*^*+/+*^ control (**Fig.1E, SFig. 1A, STable 1**,**2**). The overlap between DEGs at the two developmental time points was 64% (**SFig. 1B**).

Further, to differentiate changes driven by nuclear depletion from those caused by the expression of mutant FUS, we performed RNA-seq on 3 month-old *Fus* knockout (*Fus*^*-/-*^) animals and littermate controls, which are also viable from C57BL/6J × DBA/2J intercrossing. Differential expression analysis revealed widespread changes in gene expression, with 1,236 DEGs in *Fus*^*-/-*^ spinal cords compared to controls (**Fig. 1F, STable 3**). Both Δ14FUS expression and loss of FUS led to alterations in FUS-binding transcripts ^22^, as well as non-target RNAs (**Fig. 1E,F, SFig. 1C**). Transcripts encoding TAF15 and EWSR1, which together with FUS form the FET RBP family, were among the directly bound RNAs and showed increased expression, albeit to a different extent (**STable 2**,**3**; *Taf15* log_2_ fold change in *Fus*^*Δ14/Δ14*^ = 0.47 ± 0.06, *Fus*^*-/-*^ = 0.5 ± 0.06; *Ewsr1* log_2_ fold change in *Fus*^*Δ14/Δ14*^ = 0.17 ± 0.03, *Fus*^*-/-*^ = 0.16 ± 0.04; *P* adj < 10^−4^).

Overall, 85% of the altered transcripts in *Fus*^*Δ14/Δ14*^ mice overlapped with those in the *Fus*^*-/-*^ dataset, with a significant correlation observed between changes (**Fig. 1G,H**; Pearson correlation coefficient R^2^ = 0.4, *P* < 2 × 10^−16^), supporting a loss of function mechanism in *Fus*^*Δ14/Δ14*^ adult mutants. Interestingly, gene ontology (GO) analysis of the 1 month dataset mostly highlighted terms related to ‘lipid metabolism’, whereas the 3 month analysis identified ‘translation’, ‘ribosomal complex’ and ‘synaptic’ as enriched terms in both *Fus*^*Δ14/Δ14*^ and *Fus*^*-/-*^ datasets (**SFig. 1D-F**), possibly reflecting earlier and later ALS phenotypes, respectively.

### 2. Splicing alterations, including RBP intron retention events, contribute to FUS loss of function in adult *Fus*^*Δ14/Δ14*^ mice

In addition to transcriptional regulation, FUS regulates splicing. Splicing analysis revealed a correlation in overall splicing changes between *Fus*^*Δ14/Δ14*^ and *Fus*^*-/-*^ datasets, except for three non-concordant events (*Prune2, Exoc6b* and *Mtmr14*) (**SFig. 1G-I, STable 4**; Pearson correlation coefficient R^2^ = 0.54, *P* = 2 × 10^−5^). GO term analysis of splicing alterations highlighted terms related to ‘actin cytoskeleton’ for the *Fus*^*Δ14/Δ14*^ dataset, and ‘synaptic regulation’, ‘microtubule anchoring’ and ‘neurogenesis’ for the *Fus*^*-/-*^ dataset (**SFig. 1J,K**).

We have previously shown that FUS undergoes autoregulation through an intron retention (IR) event, a mechanism impaired by FUS nuclear depletion ^23^. We assessed this event in adult *Fus*^*Δ14/Δ14*^ mice and found an overt decrease in intron 6 and 7 reads in *Fus*^*Δ14/Δ14*^ spinal cord samples compared to *Fus*^*+/+*^ controls (**Fig. 1I**; *Fus* IR ratio in *Fus*^*+/+*^ = 0.12 ± 0.02, *Fus*^*Δ14/Δ14*^ 0.02 ± 0.003). Consistently, we found that mature *Fus* RNA (containing no intron reads) and protein levels were increased in mutant spinal cords (**Fig. 1J, SFig. 1L,M**; normalised *Fus* reads in *Fus*^*+/+*^ = 4,938.58 ± 167.43, *Fus*^*Δ14/Δ14*^ = 5,744.69 ± 97.71, *n* = 4-5; FUS protein levels in *Fus*^*+/+*^= 0.34 ± 0.09, *Fus*^*+/Δ14*^= 0.96 ± 0.19, *Fus*^*Δ14/Δ14*^= 2.24 ±

0.53, *n* = 3), confirming that FUS autoregulation is impaired in adult FUS-ALS mice.

Interestingly, *Taf15* and *Ewsr1* also showed decreased IR in spinal cord samples from *Fus*^*Δ14/Δ14*^ and *Fus*^*-/-*^ mice compared to *Fus*^*+/*+^ controls (**SFig. 1N,O**; *Taf15* IR ratio for intron 7 in *Fus*^*+/+*^ = 0.14 ± 0.01, *Fus*^*Δ14/Δ14*^ 0.08 ± 0.002; *Ewsr1* IR ratio for intron 8 in *Fus*^*+/+*^ = 0.32 ± 0.02, *Fus*^*Δ14/Δ14*^ 0.18 ± 0.01; *Ewsr1* IR ratio for intron 9 in *Fus*^*+/+*^ = 0.25 ± 0.02, *Fus*^*Δ14/Δ14*^ 0.12 ± 0.006), suggesting possible cross-regulatory mechanisms among FET proteins through IR.

### 3. Translatome alterations reveal an *in vivo* cytoplasmic gain of function in adult *Fus*^*Δ14/Δ14*^ mice

Increased cytoplasmic localisation of FUS-ALS mutants leads to cytoplasmic gain of function mechanisms, including the aberrant phase-separation of FMRP, a process associated with repressed translation of FMRP-bound RNAs ^10^. We investigated the impact of FUS mislocalisation on FMRP partitioning in adult *Fus*^*Δ14/Δ14*^ mice, and found that FMRP puncta density was significantly increased in *Fus*^*Δ14/Δ14*^ spinal motor neurons (**Fig. 2A,B**; average puncta density for *Fus*^*+/+*^ = 20.54 ± 7.03, *Fus*^*Δ14/Δ14*^ = 42.13 ± 2.93, *n* = 6). Next, we investigated whether the aberrant partitioning correlated with translatome alterations in adult *Fus*^*Δ14/Δ14*^ mice. To profile the motor neuron transcriptome, we generated triple transgenic animals expressing the Cre-dependent hemagglutinin (HA)– tagged Rpl22 ribosomal subunit (Rpl22^HA^, RiboTag), a motor neuron-specific Cre-recombinase (Chat-Cre), and either *Fus*^*+/+*^ or *Fus*^*Δ14/Δ14*^, to obtain *Fus*^*+/+*^*/Chat-cre/Rpl22*^*HA*^ and *Fus*^*Δ14/Δ14*^*/Chatcre/Rpl22*^*HA*^ animals. This strategy allowed the selective immuno-purification of ribosome-bound RNAs (translatome hereafter) from spinal cord motor neurons (**Fig. 2C**). Translatome analysis revealed global alterations in *Fus*^*Δ14/Δ14*^ compared to *Fus*^*+/+*^ datasets (**Fig. 2D**, STable 5; 96 upregulated transcripts, 62 downregulated transcripts, *n* = 5).

**Fig. 2.**
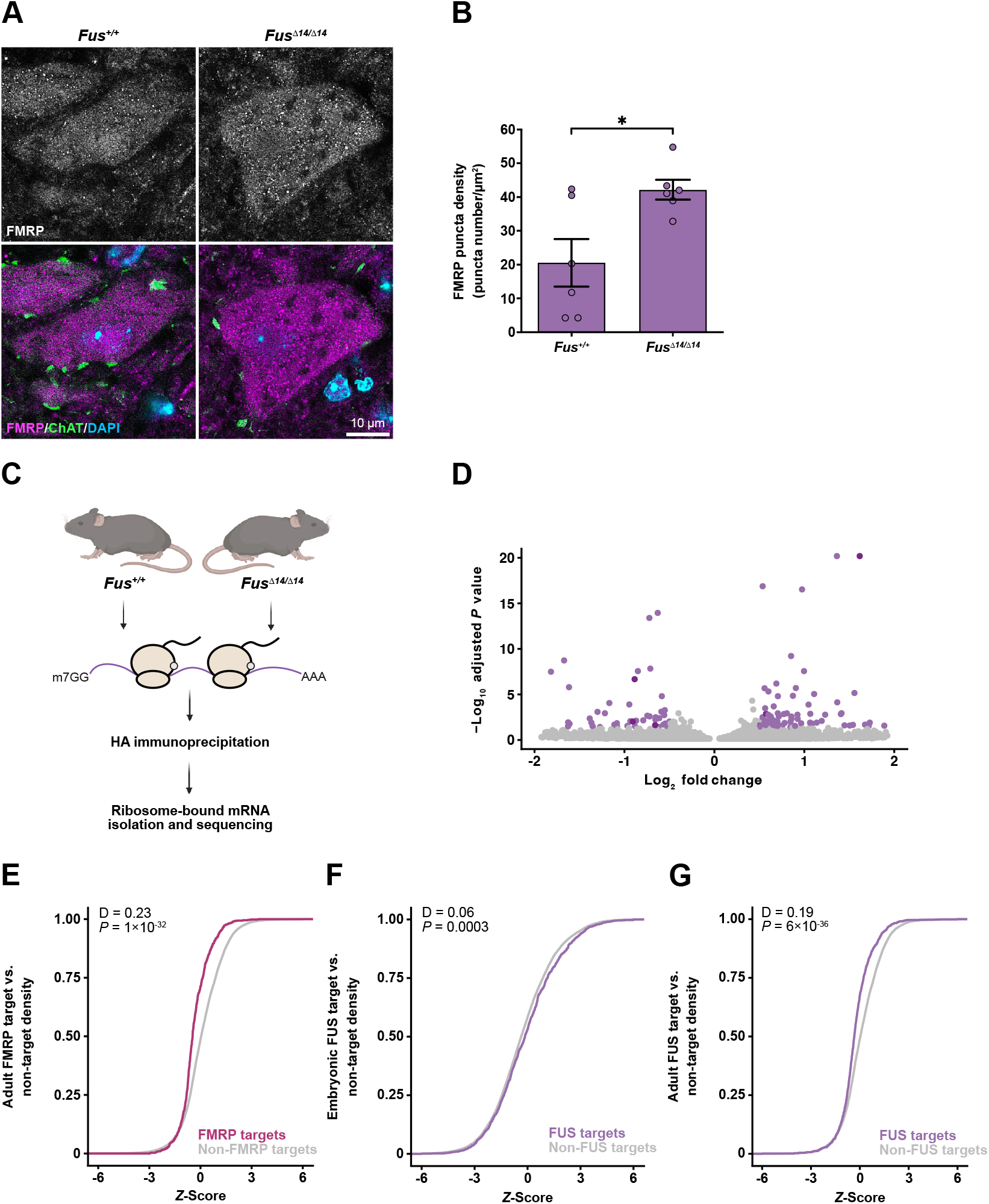
Δ14FUS expression leads to FMRP partitioning and translational alterations. (**A**) Representative single plane images of *Fus*^*+/+*^ (left) and *Fus*^*Δ14/Δ14*^ (right) motor neurons in lumbar (L)1-L6 spinal cord. FMRP is depicted in magenta, cholinergic neurons (ChAT) in green, and nuclei (DAPI) in blue. Scale bar = 10 µm. (**B**) FMRP puncta density is significantly increased in motor neurons from *Fus*^*Δ14/Δ14*^ mice compared to *Fus*^*+/+*^ controls (*P* = 0.018, unpaired *t*-test). *n* = 6. (**C**) RiboTag was used to purify motor neuron-specific, ribosome-associated transcripts from spinal cords of adult *Fus*^*+/+*^ and *Fus*^*Δ14/Δ14*^ mice. (**D**) Volcano plot showing translatome changes in *Fus*^*Δ14/Δ14*^ versus *Fus*^*+/+*^ datasets at 3 months. Known FUS targets are presented as darker dots, and non-significant changes (adj. *P* ≥ 0.05) are depicted in grey. (**E**) Cumulative frequency plot of *Z*-scores show a significant decrease of FMRP targets (plum) compared to non-target RNAs (grey) with similar expression levels in ribosomal fractions (translatome) of *Fus*^*Δ14/Δ14*^ versus *Fus*^*+/+*^ adult mice (*P* = 1 × 10^−32^, Kolmogorov-Smirnov). (**F**) Cumulative frequency plot of *Z-*scores show a slight increase of FUS targets (purple) compared to non-target RNAs (grey) with similar expression levels in ribosomal fractions in embryonic (E17.5) translatome of *Fus*^*Δ14/Δ14*^ versus *Fus*^*+/+*^ mice (*P* = 0.0003, Kolmogorov-Smirnov). (**G**) Cumulative frequency plot of *Z-*scores show a significant decrease of FUS targets (purple) compared to non-target RNAs (grey) with similar expression levels in ribosomal fractions of *Fus*^*Δ14/Δ14*^ versus *Fus*^*+/+*^ adult mice (P = 6 × 10^−36^, Kolmogorov-Smirnov).

To look specifically at FMRP target RNAs in this dataset, we plotted the cumulative density of their *Z*-scores compared to those of non-target RNAs expressed at similar levels. In this analysis, a rightward shift of the cumulative distribution curve represents an increased ribosome-RNA association, whereas a leftward-shift indicates a decreased association. We found that FMRP-bound RNAs were significantly depleted from ribosomal immuno-purified fractions compared to non FMRP-bound RNAs in spinal motor neurons from *Fus*^*Δ14/Δ14*^ mice versus *Fus*^*+/+*^ controls (**Fig. 2E**; D = 0.23, *P* = 1 × 10^−32^). This depletion in ribosome-associated FMRP targets was greater compared to the change in their total expression levels (**SFig. 2A**; D= 0.16, *P* = 7.8 × 10^−16^), corroborating our *in vivo* E17.5 embryonic datasets ^10^. While the E17.5 translatome dataset showed a slight increase in both the expression and ribosomal association of FUS-bound mature RNAs (presenting FUS binding in the 5′ UTR, open reading frame and 3′ UTR) (**Fig. 2F, SFig. 2B**; translatome D = 0.06, *P* = 0.0003, total D = 0.13, *P* = 3.2 × 10^−16^), in adult mice, FUS targets were decreased in *Fus*^*Δ14/Δ14*^ versus *Fus*^*+/+*^ ribosomal fractions compared to non FUS-bound RNAs with similar expression levels (**Fig. 2G**; D = 0.19, *P* = 6 × 10^−36^), despite a minor increase in their total expression (**SFig. 2C**; D = 0.12, *P* = 1.4 × 10^−13^). These data highlight that mutant FUS triggers translational deficits in adult *Fus*^*Δ14/Δ14*^ animals *in vivo*, with a stronger impact on FUS-bound RNAs compared to earlier developmental stages.

### 4. Neuromuscular junction innervation is impaired in *Fus*^*Δ14/Δ14*^ females

Axonal degeneration and NMJ denervation are early pathological features of ALS and precede motor neuron loss ^24,25^. We assessed the impact of Δ14FUS expression on the adult neuromuscular system by analysing NMJ innervation in fast-twitch hind-limb lumbrical muscles, which enable whole muscle analysis devoid of sectioning artefacts (**Fig. 3A**). NMJs were scored as fully innervated (∼ 80-100% overlap between axonal/presynaptic and postsynaptic compartments), partially innervated (< 80%) or denervated (no axonal/presynaptic input) (**Fig. 3A**). Interestingly, NMJ innervation in *Fus*^*Δ14/Δ14*^ animals clustered into two distinct populations. Subsequent sex separation revealed that female *Fus*^*Δ14/Δ14*^ mice, but not males, exhibited a reduced percentage of fully innervated NMJs, and a proportional increase in partially innervated NMJs, while denervated NMJs were unaffected (**Fig. 3B, STable 6**). While no overt alterations in NMJ innervation were present in male *Fus*^*Δ14/Δ14*^ mice, both male and female *Fus*^*Δ14/Δ14*^ mutants displayed significantly reduced endplate size (**Fig. 3C, SFig. 3A, STable 6**), consistent with a FUS-ALS postsynaptic muscle phenotype ^26^. No evidence of motor neuron loss was observed at this time point between *Fus*^*Δ14/Δ14*^ mice and *Fus*^*+/+*^ controls (**SFig. 3B, STable 6**).

**Fig. 3.**
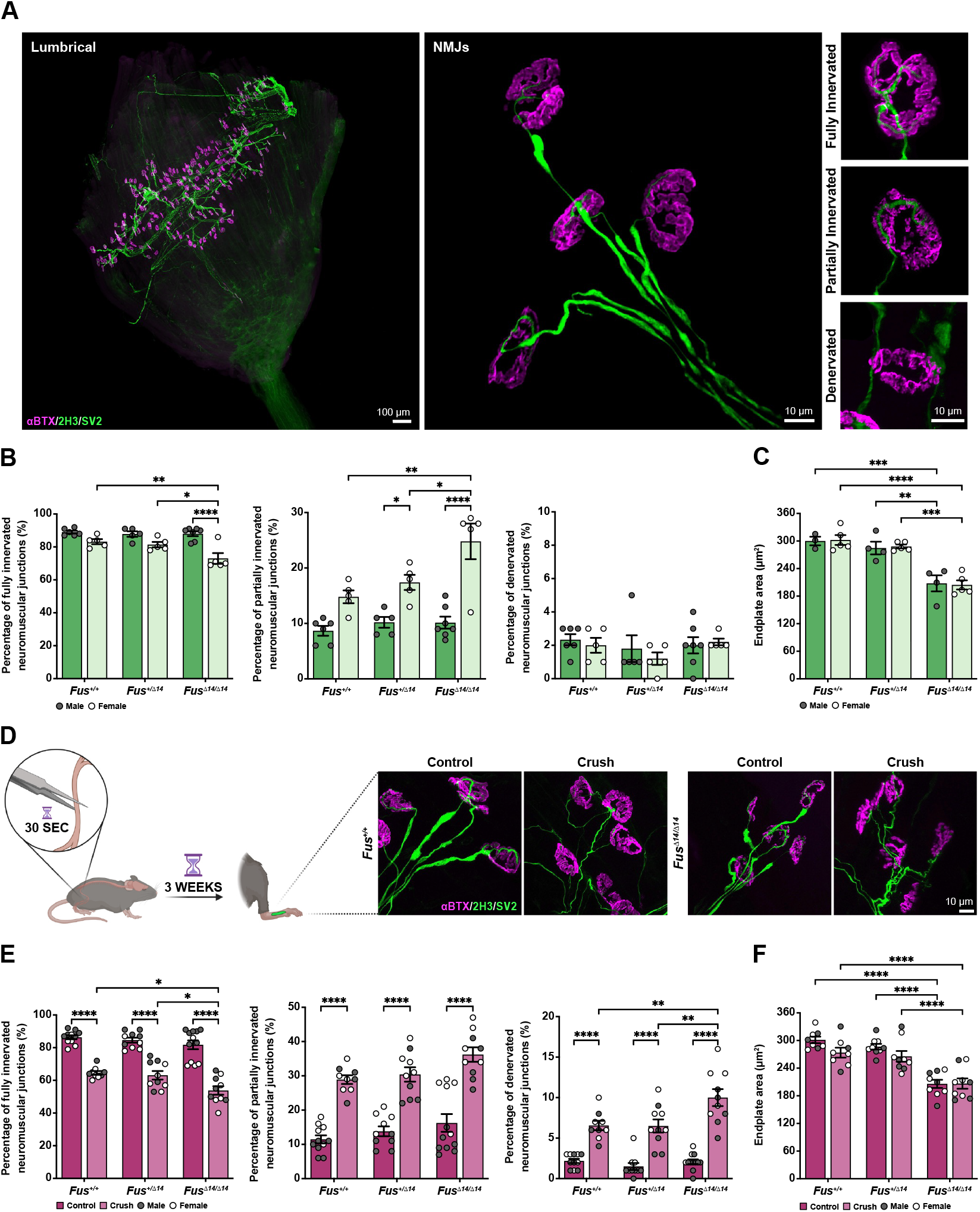
Female *Fus*^*Δ14/Δ14*^ mice display impaired neuromuscular junction innervation, whereas injury-driven reinnervation is equally disrupted in both sexes. (**A**) Representative maximum z-stack projection images of neuromuscular junctions (NMJs) in hind-paw lumbrical muscles. The entire neuromuscular innervation pattern can be observed following wholemount immunofluorescence (left). Individual NMJs can be assessed to determine innervation status (centre), and are categorised as fully innervated (80 - 100% overlap between the axonal/presynaptic and postsynaptic components), partially innervated (< 80% overlap) or denervated (no overlap) (right). Synaptic vesicles and neurofilament M (green) are co-visualised using antibodies against SV2 and 2H3, allowing simultaneous labelling of the presynaptic and axonal regions, respectively. Postsynaptic acetylcholine receptors (AChRs) are labelled with αBTX (magenta). Scale bars = 100 µm for lumbrical; 10 µm for NMJs. (**B**) Female *Fus*^*Δ14/Δ14*^ mice present a significant reduction in the percentage of fully innervated NMJs (*P* = 0.01 for genotype, *P* < 0.0001 for sex, *P* = 0.03 for interaction; two-way ANOVA), and a concomitant increase in the percentage of partially innervated NMJs (*P* = 0.003 for genotype, *P* < 0.0001 for sex, *P* = 0.02 for interaction; two-way ANOVA). This innervation deficit is not observed in male mutants. Levels of denervation are similar between sexes and across genotypes (*P* = 0.34 for genotype, *P* = 0.54 for sex, *P* = 0.70 for interaction; two-way ANOVA). *n* = 5-7. (**C**) *Fus*^*Δ14/Δ14*^ mice display significantly reduced endplate size compared to *Fus*^*+/Δ14*^ and *Fus*^*+/+*^ animals, independent of sex (*P* < 0.0001 for genotype, *P* = 0.94 for sex, *P* = 0.95 for interaction; two-way ANOVA). *n* = 3-5. (**D**) Schematic of the sciatic nerve crush paradigm. External pressure was applied to the exposed nerve for 30 sec, and lumbrical NMJ reinnervation analysed 3 weeks after crush. Representative maximum z-stack projection images depict NMJs in *Fus*^*+/+*^ (left) and *Fus*^*Δ14/Δ14*^ (right) mice under control and crush conditions. Postsynaptic AChRs were labelled with αBTX (magenta), and anti-SV2/2H3 applied for visualisation of the presynaptic/axonal components (green). Scale bar = 10 µm. (**E**) Following sciatic nerve crush, *Fus*^*Δ14/Δ14*^ mice present significantly reduced levels of fully innervated NMJs compared to *Fus*^*+/Δ14*^ and *Fus*^*+/+*^ animals (*P* = 0.0015 for genotype, *P* < 0.0001 for experimental condition, *P* = 0.24 for interaction; two-way ANOVA), and a concomitant increase in the percentage of denervated NMJs (*P* = 0.003 for genotype, *P* < 0.0001 for experimental condition, *P* = 0.01 for interaction; two-way ANOVA). Males and females respond similarly to injury. *n* = 9-12. (**F**) Sciatic nerve crush does not alter endplate area in any genotype (*P* < 0.0001 for genotype, *P* = 0.046 for experimental condition, *P* = 0.31 for interaction; two-way ANOVA). *n* = 8-9. For all graphs, \**P* < 0.05, \*\**P* < 0.01, \*\*\**P* < 0.001, \*\*\*\**P* < 0.0001 with Bonferroni’s multiple comparisons. Grey circles = males, white circles = females.

### 5. Axonal regeneration is impaired in both *Fus*^*Δ14/Δ14*^ females and males

To further investigate the Δ14FUS axonal phenotype *in vivo*, we examined NMJ reinnervation following axonal damage. We performed unilateral sciatic nerve crush and assessed NMJ reinnervation in lumbrical muscles compared to contralateral, non-injured control 3 weeks post-surgery (**Fig. 3D, SFig. 3C**). Analysis of *Fus*^*+/+*^ animals revealed that ∼ 65% of NMJs were fully reinnervated, ∼ 29% were only partially reinnervated and ∼ 7% remained denervated. While no significant difference was observed between *Fus*^*+/+*^ and *Fus*^*+/Δ14*^ animals, *Fus*^*Δ14/Δ14*^ males and females showed a significant decrease in NMJ reinnervation compared to *Fus*^*+/+*^ and *Fus*^*+/Δ14*^ mice (**Fig. 3E, STable 6**). Endplate area was not altered following crush and was comparable to contralateral, non-crush muscles of each genotype (**Fig. 3F, STable 6**).

### 6. Δ14FUS expression results in deficits in the axonal transport of mitochondria *in vivo*

Axonal transport deficits occur pre-symptomatically in several ALS models ^27^. Among these, mitochondrial dynamics play a critical role in the maintenance of axonal homeostasis and are dysregulated in ALS models ^28^. We therefore investigated *in vivo* mitochondrial transport dynamics in *Fus*^*Δ14/Δ14*^ mutants and their littermate controls. By crossing *Fus*^*+/Δ14*^ animals with transgenic mice expressing a mitochondrially targeted cyan fluorescent protein (CFP hereafter mitoCFP) in a subset of neurons that included spinal motor neurons (mitoCFP-S ^29^), we obtained adult *Fus*^*+/+*^*/mitoCFP* and *Fus*^*Δ14/Δ14*^*/mitoCFP* mice. We performed intramuscular injections of a fluorescent retrograde tracer to selectively image mitochondria within axons innervating the fast-twitch *tibialis anterior* (TA) muscle, which shows early pathological deficits in FUS and SOD1 models of ALS ^18,30^. Both average and maximum mitochondria speeds were significantly reduced in anterograde and retrograde directions in *Fus*^*Δ14/Δ14*^ mice compared to *Fus*^*+/+*^ controls (**Fig. 4A-C, SFig. 4A-C, SVideo 1, STable 6**). This decrease in speed was associated with increased pausing (**Fig. 4D, STable 6**). While no differences were observed in average or maximum speeds between sexes (**SFig. 4D-G, STable 6**), pausing was significantly increased in *Fus*^*Δ14/Δ14*^ females, specifically in anterogradely moving mitochondria (SFig. 4H,I, **STable 6**). Impaired transport dynamics were not associated with alterations in mitochondrial area, occupancy or density in *Fus*^*Δ14/Δ14*^ mice compared to *Fus*^*+/+*^ controls (**Fig. 4E,F, SFig. 4J**).

**Fig. 4.**
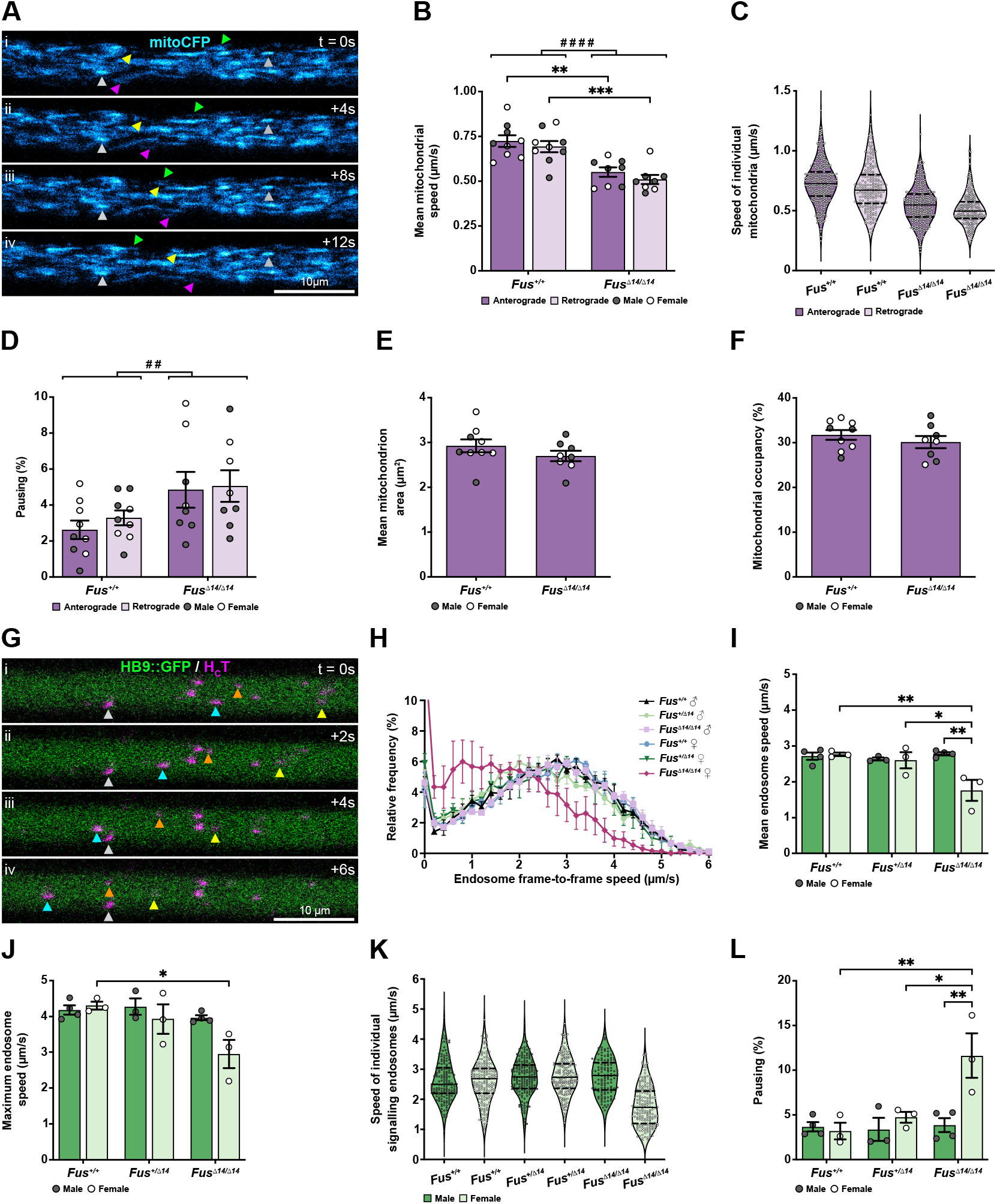
*In vivo* mitochondrial transport is impaired in *Fus*^*Δ14/Δ14*^ mice, whereas axonal transport of signalling endosomes is only affected in *Fus*^*Δ14/Δ14*^ females. (**A**) Representative time-lapse images of mitoCFP-positive mitochondria within a 12 s interval, recorded from a sciatic nerve axon *in vivo*. Grey arrowheads show stationary mitochondria, yellow and pink arrowheads label anterogradely moving mitochondria, and green arrowheads indicate a retrogradely moving mitochondrion. Scale bar = 10 µm. (**B**) Mitochondrial transport analysis reveals slower mean speeds of anterogradely and retrogradely moving mitochondria in *Fus*^*Δ14/Δ14*^ mice compared to *Fus*^*+/+*^ controls (*P* < 0.0001 for genotype, *P* = 0.23 for direction, *P* = 0.86 for interaction; two-way ANOVA). *n* = 8-9. (**C**) Violin plots show the distribution of individual mitochondrial speeds (*Fus*^*+/+*^ anterograde = 386, retrograde = 210; *Fus*^*Δ14/Δ14*^ anterograde = 365, retrograde = 233). (**D**) Mitochondria spend a greater percentage of time paused in *Fus*^*Δ14/Δ14*^ mice compared to *Fus*^*+/+*^ controls (*P* = 0.009 for genotype, *P* = 0.55 for direction, *P* = 0.75 for interaction; two-way ANOVA). *n* = 8-9. There are no alterations in (**E**) mean mitochondrion area (*P* = 0.25, unpaired *t*-test) or (**F**) occupancy (*P* = 0.37, unpaired *t*-test). *n* = 8-9. (**G**) Representative time-lapse images of H^C^T-positive signalling endosomes (magenta) within a 6 s interval, recorded from a sciatic nerve axon defined by GFP expression *in vivo* (green). Colour-coded arrowheads (cyan, orange and yellow) identify three endosomes undergoing active retrograde transport, whereas grey arrowheads indicate a stationary endosome. Scale bar = 10 µm. (**H**) Speed frequency curves show a decrease in signalling endosome frame-to-frame speeds in *Fus*^*Δ14/Δ14*^ females. Speed analysis highlights slower (**I**) mean speed (*P* = 0.015 for sex, *P* = 0.015 for genotype, *P* = 0.004 for interaction; two-way ANOVA) and (**J**) maximum speed (*P* = 0.053 for sex, *P* = 0.01 for genotype, *P* = 0.08 for interaction; two-way ANOVA) in *Fus*^*Δ14/Δ14*^ females. *n* = 3-4. (**K**) Violin plots show the distribution of individual signalling endosome speeds (*Fus*^*+/+*^ males = 168, females = 188; *Fus*^*+/Δ14*^ males = 187, females = 189; *Fus*^*Δ14/Δ14*^ males = 126, females = 187). (**L**) Signalling endosomes in *Fus*^*Δ14/Δ14*^ females spend a greater percentage of time paused (*P* = 0.01 for sex, *P* = 0.005 for genotype, *P* = 0.009 for interaction; two-way ANOVA). *n* = 3-4. For all graphs, \**P* < 0.05, \*\**P* < 0.01, \*\*\**P* < 0.001 with Bonferroni’s or Šídák’s multiple comparisons. # shows comparisons between genotypes. Grey circles = males, white circles = females.

*7. In vivo* transport of signalling endosomes is selectively impaired in *Fus*^*Δ14/Δ14*^ females

Signalling endosomes are retrogradely transported organelles that mediate long distance neurotrophic signalling, a process that is dysfunctional in pre-symptomatic ALS mouse models ^31^. We investigated whether the transport of these organelles was affected in *Fus*^*Δ14/Δ14*^ mutants. To specifically select motor axons within the sciatic nerve, we crossed mice expressing GFP under the motor neuron promoter HB9 (HB9::GFP) with *Fus*^*+/Δ14*^ mice to generate adult *Fus*^*+/+*^*/HB9::GFP, Fus*^*+/Δ14*^*/HB9::GFP* and *Fus*^*Δ14/Δ14*^*/HB9::GFP* animals. Signalling endosomes were visualised by injecting the TA with a fluorescently-labelled atoxic heavy chain domain of the tetanus neurotoxin (H_C_T), which is retrogradely transported within neurotrophin-containing signalling endosomes from axonal terminals to neuronal cell bodies ^32^ (**Fig. 4G; SVideo 2**). We found that signalling endosome transport speeds were reduced and pausing was increased in *Fus*^*Δ14/Δ14*^ females, but not males (**Fig. 4H-L STable 6**), a phenotype closely matching the NMJ innervation profile. Axonal caliber, measured using the GFP signal, did not differ between genotypes or sexes (**SFig. 4K, STable 6**), indicating no broad alterations in axon size in *Fus*^*Δ14/Δ14*^ mice. Collectively, these data reveal that axonal transport is impaired in *Fus*^*Δ14/Δ14*^ mutants, with signalling endosome transport impairments selectively present in mutant females, hence specifically correlating with the denervation phenotype.

## Discussion

We investigated the molecular, cellular and physiological consequences of FUS mislocalisation in adult mice *in vivo*, using a knock-in model where mutant Δ14FUS is expressed at endogenous levels. We crossed mice from two different inbred backgrounds, such that homozygous animals were viable into adulthood. This enabled the identification of ALS-associated phenotypes that were not previously observable in heterozygous mice, or that required aging up to 12-18 months ^17,18,33,34^. Δ14FUS exhibited a dose-dependent cytoplasmic enrichment, spreading throughout motor neurons and across sciatic nerve axons (**Fig. 1A-D**). We show that transcriptional, translational and mitochondrial transport alterations are present in adult male FUS-ALS mice in the absence of denervation (**Fig. 1-4**) and may predispose to pathogenesis, whereas signalling endosome transport impairments are tightly associated with denervation in female FUS-ALS mice and occur prior to motor neuron loss (**Fig. 3,4, SFig. 3B**).

Notably, while transcriptomic datasets from E17.5 embryonic *Fus*^*Δ14/Δ14*^ mice showed milder changes compared to *Fus*^*-/-*^ animals, adult *Fus*^*Δ14/Δ14*^ mutants presented more pronounced alterations. These changes were comparable to those observed in adult *Fus*^*-/-*^ mice (**Fig. 1G,H**), suggesting progression towards a more aggressive phenotype. Consistently, translatome analysis in adult *Fus*^*Δ14/Δ14*^ mice revealed a selective repression in the translation of mature FUS-bound RNAs, an effect not detected in embryonic datasets (**Fig. 2F,G**). This further supports a stronger phenotype in adult mice, in line with a predominantly adult onset disease. In our model, we did not detect significant FUS-ALS specific transcriptomic changes that were absent in FUS knockout animals ^35^. This may be due to using bulk spinal cord samples for transcriptome analysis, which could dilute motor neuron-specific alterations, or it may reflect impairments that are specific to humans.

Recent findings have identified TAF15 assembly into amyloid filaments and its localisation to FUS-positive inclusions in FTD cases ^5,36^, further highlighting a FUS-associated impairment of a broader RBP network in ALS/FTD. While we observed an upregulation of transcripts encoding the FET proteins TAF15 and EWSR1 in adult *Fus*^*Δ14/Δ14*^ mice, as previously shown in embryonic datasets ^23^, we did not observe any overt mislocalisation of the proteins in homozygous mice (data not shown). This suggests that TAF15 inclusions may require FUS aggregation, represent a late phenotype, or may result from distinct, predominantly FTD-associated alterations.

Axonal transport deficits have been reported *in vivo* at pre-symptomatic stages in several ALS models ^29–31,34^. Although both mitochondrial and signalling endosome transport are disrupted in several models of ALS ^30,34,35,37–40^, our *in vivo* results show that mitochondrial transport was affected in both male and female adult *Fus*^*Δ14/Δ14*^ mice, whereas signalling endosome transport deficits were selectively observed in female *Fus*^*Δ14/Δ14*^ mutants (**Fig. 4**), where innervation deficits were also present (**Fig. 3**). Consistent with our NMJ analysis, motor unit number estimation (MUNE) in the TA of this mouse model identified a stronger phenotype in *Fus*^*Δ14/Δ14*^ females compared to males (personal communication from Filipe Nascimento, manuscript in preparation). This corroborates the fact that long range neurotrophic signalling impairments are specifically and tightly associated with peripheral denervation in ALS, and suggests that mitochondrial transport deficits may precede or occur concurrently, but independently of signalling endosome alterations in pathogenesis. Moreover, the lack of signalling endosome transport impairments in *Fus*^*Δ14/Δ14*^ males highlights that mitochondrial transport defects are not due to a general transport deficit caused by cytoskeletal or molecular motor dysregulation. In addition, both deficits are absent in *Fus*^*Δ14/Δ14*^ primary motor neuron cultures ^10^, further supporting the fact that adult mice exhibit a more pronounced phenotype. This may reflect a progression to stronger axonal transport deficits in adult FUS-ALS mice, but could also be influenced by technical and physiological features unique to *in vivo* models, which enable the selective investigation of vulnerable axonal populations. Additionally, factors such as myelination, neuromuscular signalling, and the overall integrity of the neuromuscular system may also contribute to modulating these phenotypes ^41^.

Impaired mitochondrial transport dynamics may be associated with alterations in mitochondrial function ^38,42^, and FUS has been previously associated with mitochondrial dysfunction ^43–45^. Moreover, FUS-ALS mutants affect the expression of several mitochondrial electron transport chain components, primarily in motor neurons ^35^. Interestingly, in our mitochondrial transport analyses, although speeds were equally decreased in both *Fus*^*Δ14/Δ14*^ males and females (**Fig. 4**), females specifically presented increased pausing (**SFig. 4H,I**). This may be due to increased calcium transients associated with a hyperactivity phenotype ^46,47^, or could reflect the initial stages of a broader axonal impairment, as increased signalling endosome pausing was observed in 18 month-old *Fus*^*+/Δ14*^ mice in the absence of overall transport deficits ^34^. Mitochondrial transport deficits were not associated with morphological differences, in contrast with observations in other ALS models ^48,49^, further supporting it being an early phenotype.

Despite the difference in the baseline proportion of fully innervated NMJs between the sexes, a challenging insult, *i*.*e*., sciatic nerve crush, led to equally impaired reinnervation in *Fus*^*Δ14/Δ14*^ male and female mice (**Fig. 3**). Importantly, denervation and aberrant RBP aggregation are reversed by FUS antisense oligonucleotide treatment in FUS-ALS models ^18^, demonstrating that innervation deficits depend on mutant FUS gain of function.

Mitochondria are pivotal to sustain axonal growth and regeneration^50^. Defective mitochondrial transport in *Fus*^*Δ14/Δ14*^ animals may affect mitochondria availability and localisation to regenerating axons. Moreover, disrupted mitochondrial transport may be associated with impaired bioenergetics ^38^, which are critical for this process. Further, axonal regeneration pathways rely on fine-tuned transcription and translation ^51–53^. We and others have demonstrated that translation is impaired in FUS-ALS ^9,10,13^; in addition, FUS and FMRP target RNAs, depleted from ribosomal fractions in *Fus*^*Δ14/Δ14*^ mice, are associated with synaptic and ribosomal categories (**SFig. 1E, STable 2**), which are both critical for regrowth and innervation. Recently, Bolívar and colleagues have identified the motor neuron translational program triggered during crush-induced axonal regeneration ^51^. Interestingly, several transcripts involved in this are FUS and FMRP target RNAs (**STable 5**), and it is possible that their translation dynamics are affected following sciatic crush, highlighting that both molecular and cellular mechanisms may contribute to the reinnervation phenotype in FUS-ALS mice.

Moreover, primary embryonic motor neurons derived from *Fus*^*Δ14/Δ14*^ animals, as well as iPSC-derived *FUS*^*P525L/P525L*^ motor neurons, show increased aberrant axonal regrowth following axotomy ^11^. This could represent an early developmental phenotype, or suggest that regrowth and reinnervation *in vivo* may require both cell-autonomous and non-cell-autonomous regulated pathways, not least pathfinding, NMJ targeting and formation of stable NMJs. Altogether, our data demonstrate that adult *Fus*^*Δ14/Δ14*^ mice have progressive loss and gain of function mechanisms that result in alterations at molecular, cellular and physiological level. Our data strengthens the association between signalling endosome transport deficits and denervation in ALS, while identifying broader impairments in transport and reinnervation that may contribute to axonal vulnerability and denervation in FUS-ALS pathogenesis.

## Materials and Methods

### Animals

Animal experiments conducted in the United Kingdom adhered to the European Community Council Directive 86/609/EEC of 24 November 1986, and were performed under license from the UK Home Office in accordance with the Animals (Scientific Procedures) Act 1986 Amendment Regulations 2012. Experiments were additionally approved by the UCL Institute of Neurology Ethical Review Committee. Mice were maintained in a controlled temperature (21°C) and humidity (45-65%) environment under a 12 h light/dark cycle regime, and given *ad libitum* access to food and water.

Δ14FUS mice (B6N;B6J-Fus^tm1Emcf/H^, MGI (Mouse Genome Informatics):6100933) have been previously described ^17^. FUS knockout mice were obtained from the Mouse Knockout Project [Fustm1(KOMP)Vlcg]. Choline acetyltransferase (ChAT)–internal ribosomal entry site (IRES)–Cre [B6;129S6-*Chattm2(cre)Lowl*/J - RRID: IMSR_JAX: 006410], RiboTag (B6N.129- *Rpl22tm1*.*1Psam*/J - RRID: IMSR_JAX: 011029), mitoCFP [B6.Cg-Tg(Thy1-CFP/COX8A)S2Lich/J - RRID: IMSR_JAX: 007967], and HB9::GFP [B6.Cg-Tg(Hlxb9-GFP)1Tmj/J - RRID: IMSR_JAX: 005029] mice were obtained from the Jackson Laboratory. The frt-flanked neo cassette from B6;129S6- *Chattm2(cre)Lowl/J* mice was removed via flp-recombination to reduce ectopic expression of the transgene. Animals were backcrossed onto C57BL/6J mice for at least six generations.

To generate *Fus*^*+/Δ14*^ DBA/2J animals, *Fus*^*+/Δ14*^ mice on a congenic C57BL/6J background were backcrossed for at least ten generations on the DBA/2J genetic background. *Fus*^*+/Δ14*^ C57BL/6J females were bred with *Fus*^*+/Δ14*^ DBA/2J males to produce viable C57BL/6J-DBA/2J F1 homozygotes and control littermates. Only F1 hybrid animals were used in this study.

Adult *Fus*^*Δ14/Δ14*^*/Chat*^*cre/+*^*/Rpl22*^*HA/+*^ and *Fus*^*+/+*^*/Chat*^*cre/+*^*/Rpl22*^*HA/+*^ littermate animals were generated as follows: (1) *Chat*^*cre/+*^ mice were backcrossed 5 generations to the DBA/2J genetic background, then crossed to male *Fus*^*Δ14/+*^ (DBA2/J) mice for two subsequent generations to produce *Fus*^*Δ14/+*^*/Chat*^*cre/cre*^ (DBA/2J) mice, (2) *Rpl22*^*HA/+*^ mice were crossed to *Fus*^*Δ14/+*^ (C57BL/6J) mice for two generations to produce *Fus*^*Δ14/+*^*/Rpl22*^*HA/HA*^ (mixed C57BL/6J and C57BL/6N background) mice, (3) males from step 1 were intercrossed with females from step 2.

Adult *Fus*^*+/+*^/*mitoCFP* and *Fus*^*Δ14/Δ14*^/*mitoCFP* mice were generated by crossing *mitoCFP*^*+/-*^ C57BL/6J with *Fus*^*+/Δ14*^ C57BL/6J mice, followed by intercrossing of *Fus*^*+/Δ14*^/*mitoCFP* with DBA/2J *Fus*^*+/Δ14*^ mice.

Adult *Fus*^*+/+*^*/HB9::GFP, Fus*^*+/Δ14*^*/HB9::GFP* and *Fus*^*Δ14/Δ14*^*/HB9::GFP* mice were obtained by crossing *HB9::GFP* C57BL/6J mice with Fus^+/Δ14^ C57BL/6J mice, followed by intercrossing with DBA/2J *Fus*^*+/Δ14*^ mice.

All experiments were performed on 3 month-old mice. Preliminary analyses of neuromuscular deficits in *Fus*^*Δ14/Δ14*^ males showed ALS phenotypes, such as NMJ denervation and reduced muscle force at 3 months of age. These preliminary cohorts were generated and mostly maintained at the Harwell Institute (Oxford, UK) animal facility, whereas mice used in the remainder of this study were maintained at UCL. One key environmental difference between facilities was diet, with UCL-housed animals receiving soy-supplemented feed, which may have had an effect on hormone-regulated pathways, and as a consequence, sex-dependent ALS phenotypes observed in our cohorts ^54^.

### *In vivo* axonal transport of mitochondria and signalling endosomes

#### Intramuscular injections of HCT

A fluorescently labelled atoxic fragment of tetanus neurotoxin was prepared (H_C_T-555; residues 875–1315 fused to a cysteine-rich region and a human influenza haemagglutinin epitope) and subsequently labelled with Alexa Fluor 555 C_2_ maleimide (Thermo Fisher Scientific), as previously described ^55,56^. Pre-surgical preparations were carried out according to established protocols ^55,56^. Mice were exposed to 4-5% isoflurane (MWI Animal Health) in an induction chamber. Absence of the righting reflex was confirmed before transferring anaesthetised animals to a mouthpiece, and reducing the concentration to 2-3%. Upon absence of the pedal withdrawal reflex, a small incision was made above the upper TA muscle. 3.5 µL of 7.5-10 mg H_C_T-555 in PBS was injected using pulled, glass micropipettes (Drummond Scientific) targeting the NMJs, as previously described ^57^. The incision was sutured, and animals closely monitored for 1 h during recovery from anaesthesia.

#### In vivo imaging of axonal mitochondria and signalling endosomes

To visualise axonal mitochondrial transport, intravital imaging was performed using the mitoCFP mouse; signalling endosome transport analysis utilised the HB9::GFP mouse, as described previously ^55,56^. To identify fast α-motor neuron axons, H_C_T-555 was subcutaneously injected into the TA of *Fus*^*+/+*^/*mitoCFP* and *Fus*^*Δ14/Δ14*^/*mitoCFP* animals. Signalling endosomes were assessed in TA-innervating motor axons of *Fus*^*+/+*^*/HB9::GFP, Fus*^*+/Δ14*^*/HB9::GFP* and *Fus*^*Δ14/Δ14*^*/HB9::GFP* mice. 4-8 h post-injection, the sciatic nerve was exposed under terminal anaesthesia, and imaged on an inverted LSM780 confocal microscope (Zeiss) within an environmental chamber pre-warmed to 37°C. Time-lapse microscopy was performed using a 40×, 1.3 numerical aperture (NA) DIC Plan-Apochromat oil-immersion objective (Zeiss), with an additional 2x digital room and < 1% laser power. Nodes of Ranvier were avoided to exclude node-related changes in axonal transport dynamics ^58^. Frame intervals of 1.0-1.5 s were used to image H_C_T-555-positive signalling endosomes and 3.0-4.5 s frame intervals for mitochondria. Imaging was concluded within 1 h of initiating terminal anaesthesia.

#### Tracking analysis

Time-lapse videos were analysed in Fiji/ImageJ using the *Track-Mate* plugin ^59^. H_C_T-555-positive signalling endosomes were semi-automatically tracked, while mitochondria were tracked manually. Organelles moving for > 10 consecutive frames were included, whereas those with terminal pauses, defined as the absence of movement in > 10 consecutive frames, were excluded. A minimum of 20 trackable signalling endosomes and 10 mitochondria was analysed per axon, with > 3 individual axons assessed per mouse.

‘Mean velocity’ is defined as the overall average speed of an organelle during the entire transport event. ‘Maximum velocity’ was calculated as the average of the highest speeds of moving organelles per animal. Organelles were defined as ‘pausing’ when previously moving organelles had velocities within 0 and 0.1 µm/s between consecutive frames (to account for a potential breathing/vasculature artifact). ‘Pausing percentage’ was calculated by dividing the number of pauses by the total number of frame-to-frame movements. Mean axonal diameter was measured using the GFP signal within motor neuron axons. At least 10 orthogonal width measurements were taken per axon, and a minimum of 3 axons were analysed per animal.

### Sciatic nerve crush

Unilateral sciatic nerve crush was performed on 9 week-old mice using a procedure adapted from a previously described protocol^60^. Hind-paw lumbrical muscles from the unaffected hind-limb served as a contralateral control. Briefly, anaesthetised mice were placed in the prone position, their upper hind-limb secured at a 45° angle. Upon absence of the pedal withdrawal reflex, 250 µL of 0.5 mg/mL meloxicam (MWI Animal Health) was subcutaneously injected into the scruff. A small vertical incision was made directly below the femur, and the surrounding musculature - specifically the quadriceps femoris and biceps femoris - was separated to expose the sciatic nerve. The nerve was compressed for 30 sec, and 20 μL of 1.25 mg/mL bupivacaine (MWI Animal Health) was locally administered. The musculature was reclosed, and the incision sutured. Animals were monitored for several days post-surgery. Tissues were collected 3 weeks post-surgery (postnatal day 90).

### Dissections

Spinal cords were dissected as previously described ^61^. Hydraulic extrusion was carried out using a BD Microlance™ 3 hypodermic needle 21G x 1.5” (Becton Dickinson) attached to a 10 mL syringe containing PBS. Sciatic nerves were dissected using an adapted version of an established protocol ^62^. After exposing the sciatic nerve, perpendicular incisions were made at the most proximal and distal ends. The surrounding connective tissue was detached for complete removal of the nerve. Hind-paw lumbricals were dissected as previously described ^63,64^. Following dissection, muscles were fixed in 4% (w/v) paraformaldehyde (PFA; Thermo Fisher Scientific) in PBS for 10 min. Excess connective tissue was subsequently removed, and lumbricals stored at 4°C in 0.02% (w/v) sodium azide (Sigma) in PBS.

### Tissue processing

Sciatic nerves and spinal cords were fixed in 4% (w/v) PFA in PBS for 24 h at 4°C, washed with PBS and equilibrated in 30% (w/v) sucrose (Sigma) and 0.02% sodium azide in PBS for 24-72 h at 4°C. The lumbar(L)1–L6 region of the spinal cord was identified and isolated. Tissues were embedded in Tissue-Tek O.C.T. Compound (Sakura Finetek) and frozen. Using an OTF Cryostat (Bright Instruments), transverse sections of sciatic nerve (30 μm) and spinal cord (20 μm) were collected onto Superfrost™ Plus microscope slides (Thermo Fisher Scientific) and stored at -20°C. The spinal cord sectioning protocol for assessing motor neuron counts has been detailed elsewhere ^61^.

### Antibodies

The following primary antibodies were used: anti-ChAT [immunohistochemistry (IHC) 1:200; Sigma, AB144P], anti-FMRP [IHC 1:200; Abcam, AB17722], anti-FUS [IHC 1:300, western blot (WB) 1:5000; Novus Biologicals, NB100-565], anti-S100β [IHC 1:250; Frontier Institute, MSFR105330], anti-phospho-neurofilament H (SMI-31) [IHC 1:1000; Sigma, NE1022], anti-neurofilament M (2H3) [IHC 1:250; DSHB] and anti-SV2 [IHC 1:50; DSHB]. Alexa Fluor-conjugated secondary antibodies were purchased from Invitrogen, and used for IHC (1:500) and whole-mount immunofluorescence (1:250): donkey anti-goat 488 (A11055), donkey antimouse 488 (A21202), donkey anti-rabbit 488 (A32790), donkey anti-rabbit 555 (A32794) and donkey anti-mouse 647 (A32787). An HRP-conjugated secondary antibody was used for WB: goat anti-rabbit (1:5000; BioRad, 1706515). Alexa Fluor 555-conjugated α-bungarotoxin (1:1000; Life Technologies, B35451) was used to identify postsynaptic acetylcholine receptors (AChRs) at the NMJ, and nuclei were visualised with DAPI (1:1000; Thermo Fisher Scientific, D1306).

### Immunohistochemistry and whole-mount immunofluorescence

Heat-mediated antigen retrieval was necessary for staining ChAT in motor neurons of the spinal cord. Briefly, slides were incubated in sodium citrate buffer (pH 6.0; 10 mM tri-sodium citrate [Thermo Fisher Scientific], 0.05% Tween® 20 [Sigma]) at 90°C for 20 min, then at RT for 20 min. Slides were allowed to dry prior to IHC. Spinal cord and sciatic nerve sections were permeabilised for 10 min with 0.3% (w/v) Triton X-100 (Sigma) in PBS, then immersed in a blocking solution containing 0.5% bovine serum albumin Fraction V (Roche), 0.3% (w/v) Triton X-100 and 10% horse serum (Sigma) in PBS for 30 min. Sections were incubated overnight at 4°C with primary antibodies. Sections were then washed 3 times with PBS for 10 min, and probed with secondary antibodies for 2 h at RT. Slides were washed 3 more times with PBS for 10 min, before mounting in Fluoromount-G™ (Thermo Fisher Scientific) and concealing with 22 × 64 mm cover glass (VWR). Whole-mount immunofluorescence of lumbrical NMJs has been previously described^63,64^.

### Nissl staining

Spinal cord sections used for motor neuron counts were immersed in gallocyanin ^61^ for 30 min at RT. Slides were rinsed twice in Milli-Q water for 2 min each, before dehydrating in a series of ethanol solutions: 70% (w/v) ethanol (VWR) for 1 min, 90% (w/v) ethanol for 1 min, 100% ethanol for 2 min, then twice in Histo-Clear (National Diagnostics) for 2 min each. Slides were mounted using DPX medium (Sigma) and concealed with 22 × 64 mm cover glass.

### Confocal imaging

Imaging was performed using either an LSM 780 or LSM 980 inverted confocal microscope (Zeiss), and various objectives used: Plan-Apochromat 20×/0.8, LD LCI Plan-Apochromat 40×/1.3 Imm Corr DIC, and Plan-Apochromat 63×/1.4 Oil DIC M27 (Zeiss).

### Analyses

All analyses were performed on previously blinded samples.

#### Nuclear/cytoplasmic ratio

Nuclear and cytoplasmic FUS fluorescence intensities were quantified using Fiji/ImageJ. Nuclei were identified by DAPI signal, and cell bodies by ChAT immunolabeling. For each biological replicate, mean FUS intensity in the nucleus and cytoplasm was calculated by averaging the signal from 2 separate planes across 8-14 motor neurons.

#### FUS intensity in sciatic nerve axons

Segmentation analysis was performed using Fiji/ImageJ to quantify average FUS intensity within SMI-31 positive axons of the sciatic nerve. Z-stacks were captured at 0.5 μm intervals, and 3 focal planes selected for analysis. SMI-31 positive axons were segmented using an appropriate threshold, and smaller particles excluded using a size filter (μm^2^ = 1-infinity) under *Analyze* > *Analyze Particles*. Mean FUS intensities were averaged across slices to obtain a single measurement per image. 3 images were analysed per animal and averaged.

#### Puncta density analysis

FMRP puncta analysis was performed using Imaris (Version 9.5, Oxford Instruments). Prior to automatic granule detection, a Gaussian filter and automatic background subtraction were applied. A cell wizard layer was subsequently created to enable cellular component detection, using the ChAT channel as reference for the cell body and DAPI for the nucleus. All images were processed with identical parameters applied automatically by the software. For each image, a 3D puncta map was generated, and both puncta number and cell volume values were obtained.

#### Endplate area

Endplate area was quantified using Fiji/ImageJ, as described ^63^. For each mouse, 3 images of NMJs - each taken from different lumbrical muscles - were captured per condition, and endplate areas subsequently averaged.

#### NMJ innervation

NMJ innervation status was manually scored as previously described ^63,65^. 100 NMJs across 2 lumbrical muscles were analysed per mouse. The cut-off value for partial innervation was established as < 80%.

#### Motor neuron counts

Images of spinal cord sections were acquired using the NanoZoomer S60 Digital Slide Scanner (Hamamatsu), operating in brightfield mode at 40× magnification. Nissl-stained motor neurons in regions L3-L6, and localised to the sciatic motor pool, were counted as previously described ^61,66^.

#### Mitochondrion number, size and density analyses

Time-lapse image files with ≥ 150 original frames were analysed and batch processed in Fiji/ImageJ using a combination of plugins, including *Trainable Weka Segmentation* ^67^, *Find Stack Maxima* and *Analyze Particles*. First, equidistant frames were selected using *Slice Keeper* to create 20-frame files (e.g., using first slice “1”, last slice “220” and increment “11” would output a new 20-frame stack). Condensed stacks for each axon were individually segmented using *Trainable Weka Segmentation*, with a pre-trained machine-learning based classifier to differentiate between mitochondrial signal and background. The classifier was trained using 8 manual rounds and was able to recognize > 85% of mitochondrial signals. After segmentation, stacks of probability images were generated, providing raw masks for selecting regions with mitochondrial signal. The raw mask was improved by running *Find Stack Maxima* to identify boundary issues, *i*.*e*. several mitochondria recognised as 1 large mitochondrion. The refined mask was subsequently used to select regions with mitochondrial signal in the original 20-frame stack for analysis with *Analyze Particles*, providing total mitochondrial number and individual mitochondrion size across all 20 frames. Mitochondrial number and average mitochondrion size was calculated per axon. Mitochondria number and area were normalised to axon size to calculate mitochondria density (mitochondria number/area) and mitochondrial occupancy (mitochondria area/axonal area). At least 50 mitochondria were analysed per axon, and at least 3 individual axons assessed per mouse. Axons analysed were the same used for mitochondrial transport analysis.

### Western blotting

Adult spinal cord samples were collected via laminectomy from female mice on a C57BL/6J × DBA/2J F1 hybrid background and snap frozen. Samples were lysed in radioimmunoprecipitation assay buffer (RIPA) [50 mM tris-HCl (pH 7.5), 150 mM NaCl, 1% NP-40, 0.5% sodium deoxycholate, 0.1% SDS, 1 mM EDTA, 1 mM EGTA, Halt protease and phosphatase inhibitor cocktail (Thermo Fisher Scientific)], and incubated on a rotating wheel at 4°C for 1h. Nuclei and cellular debris were subsequently spun down at 20,000 × *g* for 15 min. Supernatants were collected, Laemmli buffer (Thermo Fisher Scientific) was added, and samples were denatured at 98°C for 5 min. Pierce™ BCA Protein Assay Kit (Thermo Fisher Scientific) was used to quantify protein concentration. Samples were electrophoretically separated using 4-15% Mini-PROTEAN® TGX Stain-Free™ protein gels (Bio-Rad). Protein content was imaged using the stain-free imaging setting in a Bio-Rad developer. Proteins were subsequently transferred onto polyvinylidene difluoride membranes through the TransBlot semidry transfer system (Bio-Rad). Membranes were blocked in 4% milk in Tris-buffered saline (Thermo Fisher Scientific) with 0.1% Tween® 20 (TSBS-T) for 1 h, followed by overnight incubation with primary antibody diluted in 4% milk in TBS-T. Membranes were washed 3 times in TBS-T and incubated with an HRP-conjugated secondary antibody for 1 h, prior to development with Immobilon Classico HRP substrate (Millipore).

### RNA extraction and sequencing

Snap frozen spinal cord samples were crushed by a pestle and mortar on dry ice, followed by lysis of half the sample in Qiazol (Qiagen). RNA was extracted using the miRNeasy Mini Kit (Qiagen) as per manufacturer’s instructions. RNA quality was checked by Tapestation (Agilent) and RNAs stored at -80°C. All sequenced samples had RIN scores higher than 8. cDNA libraries were made using the KAPA RNA HyperPrep Kit with RiboErase (Roche). Libraries were sequenced on an Illumina NovaSeq 6000 to generate paired-end 100bp reads for total RNA-seq, and on NextSeq 2000 to generate single-end 100bp for RiboTag RNA-seq.

### RiboTag immunoprecipitation

The RiboTag method was performed as described previously ^10^. Briefly, 3 month-old spinal cords were homogenized using TissueRuptor (Qiagen) in buffer containing 50 mM tris-HCl (pH 7.4), 100 mM KCl, 12 mM MgCl_2_, 1% NP-40, cycloheximide (0.1 mg/ml), heparin (1 mg/ml), and Superase·In RNase (Thermo Fisher Scientific). Lysates were cleared by centrifugation at 10,000 × *g* for 10 min, and 5% of the lysate was saved as input. To reduce non-specific binding, protein G magnetic beads (Dynabeads, Thermo Fisher Scientific) were added to the lysate and incubated for 2 h at 4°C. Next, 5 μl of anti-HA antibody (Sigma) was added to the pre-cleared lysate and incubated for 2 h at 4°C. 100 μl of beads slurry with 2 μl of Superase·In was added to the lysate, followed by 2 h incubation at 4°C. Beads were then washed 5 times in washing buffer [300 mM KCl, 1% NP-40, 50 mM tris-HCl (pH 7.4), 12 mM MgCl_2_, and cycloheximide (0.1 mg/ml)]. Beads were eluted in Qiazol, and RNAs isolated using the RNeasy Micro Kit (Qiagen). The quality control of RNA was performed using TapeStation (Agilent).

### RNA-seq and splicing analysis

Raw sequences (in FASTQ format) were trimmed using Fastp ^68^ with the parameter *qualified_quality_phred: 10*, and aligned using STAR (v2.7.0f) ^69^ to the *Mus musculus* genome assembly GRCm38. Counting the reads was performed using featureCounts v1.6.4 ^70^. Differential gene expression was performed using DESeq2 ^71^. The significance level was set at an adjusted *P* value of 0.05. IR events were measured using IRFinder v1.3.1 ^72^ with default settings in the BAM mode. Splicing analysis was performed using MAJIQ (v2.1) ^73^ on sorted BAMs and the GRCm38 reference genome. MAJIQ’s outputs with probability_changing > 0.9 and probability_non_changing < 0.05 were used to categorise each of the splicing junctions by the overlap with annotated transcripts and exons ^74^, using GTF and DASPER (https://github.com/dzhang32/dasper). For certain alternative splicing events, there was only enough coverage to call one of the two defining junctions significant. To perform translatome analysis, each nominal *P* value was converted into a *Z*-score and given the sign of the log_2_ fold change. A list of FMRP target genes was obtained from a published HITS (high-throughput sequencing)-CLIP experiment ^75^. A list of FUS target peaks was obtained from a published iCLIP experiment ^22^. For translatome analysis, FUS targets were filtered to only use peaks overlapping coding and UTR regions. Non-target transcripts with base mean (log_10_) > 0.75 and base mean (log_10_) < 3.25 were used for comparison. GO terms analysis was performed in R using the enrichGO function from the clusterProfiler package ^76^.

### Statistical analysis

Data normality was evaluated using the Shapiro-Wilk test, and homogeneity of variance was presumed. Unpaired *t*-tests, one-way analysis of variance (ANOVA) and two-way ANOVA were used for statistical comparison of parametric datasets, with the latter 2 assessments incorporating Bonferroni’s or Šídák’s multiple comparisons test. Equivalent non-parametric alternatives were considered when data deviated from standard assumptions. In such cases, datasets were analysed using the Kruskal-Wallis test with Dunn’s multiple comparisons. Statistical analyses were two-tailed, with significance established at an α-level of *P* < 0.05. Statistical computations and graph generation were performed using GraphPad Prism 10 (Version 10.4.1; San Diego, California). Sample sizes, pre-determined using power calculations, are reported in figure legends and represent biological replicates. Data are presented as mean ± SEM.

## Acknowledgements

We thank staff of the Denny Brown Laboratory at the Queen Square Institute of Neurology for looking after our colonies at UCL, in particular Catherine Hills and Zoe Windsor. We thank staff involved at the Mary Lyon Centre at MRC Harwell, including Lydia Teboul and the Molecular and Cellular Biology Team for genotyping; Jackie Harrison, Gemma Atkins, Dr Anju Paudyal, Emily Ireson, and Dr Michelle Stewart for animal husbandry and colony management. We thank Filipe Nascimento and all members of the Fratta, Schiavo and Sleigh Laboratories for critical discussions. Schematics in Figure 2C and 3D were generated using BioRender (https://www.biorender.com/).

This work was supported by the following awards:

UK Medical Research Council Senior Clinical Fellowship MR/M008606/1 (PF); Motor Neurone Disease Association Lady Edith Wolfson Fellowship MR/S006508/1 (PF). The Motor Neuron Disease Association fellowships: Birsa/Oct21/976-799 (NB) and Tosolini/Oct20/973-799 (APT). Col Bambrick MND Research Grant from Motor Neuron Disease Research Australia (IG 2450) (APT); FightMND Drug Development Grant awarded to Giovanni Nardo (Istituto di Ricerche Farmacologiche Mario Negri - IRCCS) (DDG-73; for APT). Wellcome Trust Senior Investigator Awards (107116/Z/15/Z and 223022/Z/21/Z) (GS), and a UK Dementia Research Institute award (UKDRI-1005) (GS). The UK Medical Research Council (MRC) awards: MC_EX_MR/N501931/1 (EMCF) and MR/Y010949/1 (JNS). The UCL Therapeutic Acceleration Support scheme supported by funding from MRC IAA 2021 UCL MR/X502984/1 (JNS); the China Scholarship Council (QL).

## Author contribution

Conceptualisation: PF, NB, APT, GS, BK

Data Curation and Formal analysis: AM, WJ, QL, FL

Investigation: RLS, APT, AM, AMU, WJ, BK, MA, MFM, DVC, NB, TJC

Resources: MFM, GP, TJC

Writing - Original draft: NB, APT, RLS

Writing - Review and editing: all authors

Supervision: NB, PF, LG, JNS, GS

Funding Acquisition: PF, NB, APT, JNS, GS, EMCF

## Data Availability

All data needed to evaluate the conclusions in the paper are present in the paper and/or in the Supplementary Materials. Raw RNA-sequencing data were deposited on Gene Expression Omnibus (GEO) with accession numbers: GSE298456 (RNA-Seq) and GSE298457 (RiboTag).

## Supplementary Figures

**Fig. S1.**
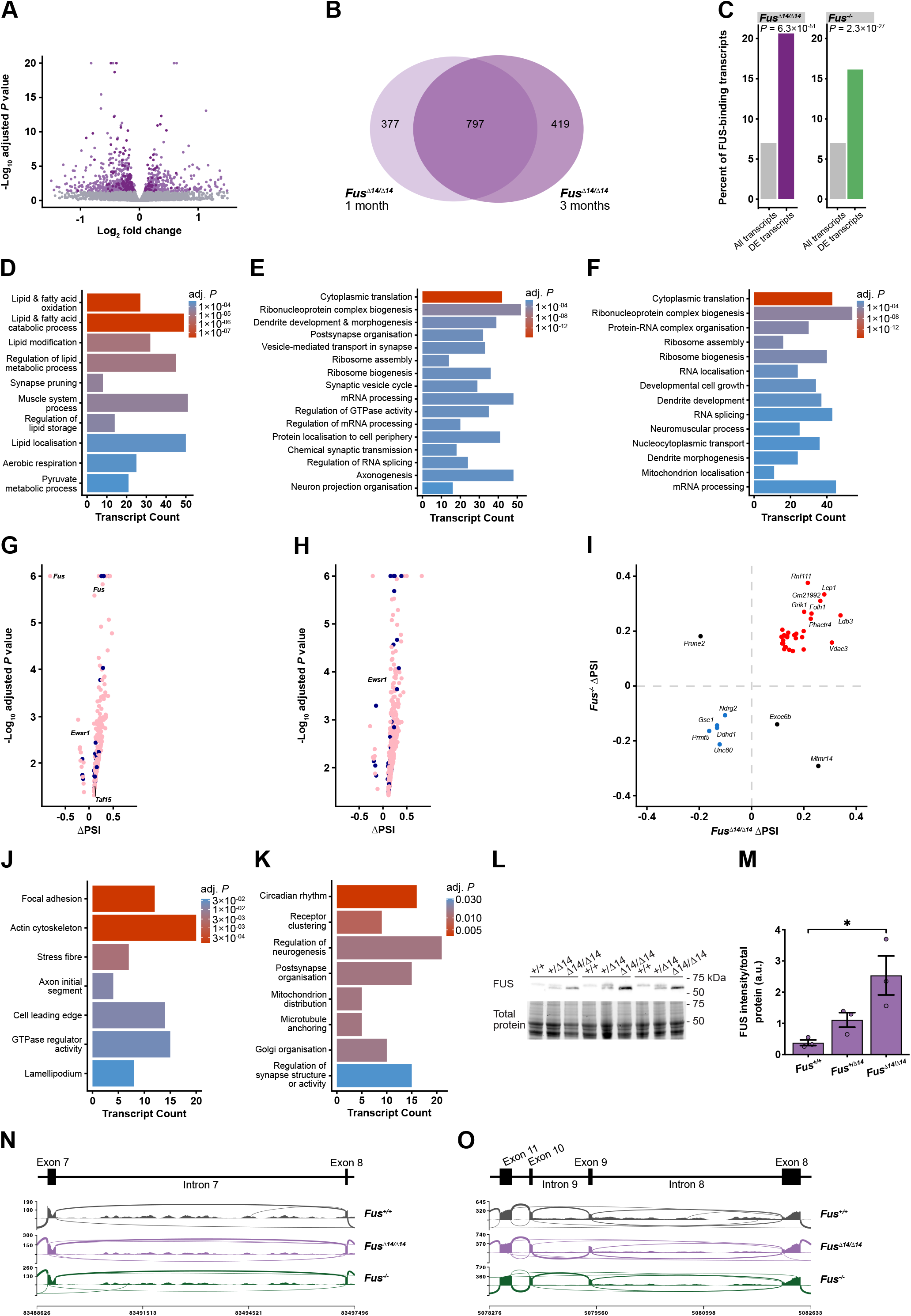
Δ14FUS expression results in transcriptional and splicing alterations consistent with nuclear loss of function in adult mice. (**A**) Volcano plot showing differentially expressed genes in *Fus*^*Δ14/Δ14*^ versus *Fus*^*+/+*^ datasets at 1 month. Known FUS targets are presented as darker dots, and non-significant changes (adj. *P* ≥ 0.05) are depicted in grey. (**B**) Venn diagram showing the overlap between differentially expressed genes identified in *Fus*^*Δ14/Δ14*^ versus *Fus*^*+/+*^ at 1 month (light purple) and *Fus*^*Δ14/Δ14*^ versus *Fus*^*+/+*^ datasets at 3 months (dark purple). (**C**) FUS-bound transcripts are significantly enriched among differentially expressed genes compared to the whole transcriptome in *Fus*^*Δ14/Δ14*^ and *Fus*^*-/-*^ datasets (20.7% versus 7%, *P* = 6.3 × 10^−51^ in *Fus*^*Δ14/Δ14*^ versus *Fus*^*+/+*^; 16.1% versus 7%, *P* = 2.3 × 10^−27^ in *Fus*^*-/-*^ versus *Fus*^*+/+*^). (**D**) Gene Ontology (GO) enrichment analysis of differentially expressed genes identified in *Fus*^*Δ14/Δ14*^ versus *Fus*^*+/+*^ datasets at 1 month. The color intensity corresponds to the *P* value. (**E**) GO enrichment analysis of differentially expressed genes identified in *Fus*^*Δ14/Δ14*^ versus *Fus*^*+/+*^ datasets at 3 months. (**F**) GO enrichment analysis of differentially expressed genes identified in *Fus*^*-/-*^ versus *Fus*^*+/+*^ datasets at 3 months. (**G**) Volcano plot showing splicing alterations in *Fus*^*Δ14/Δ14*^ versus *Fus*^*+/+*^ datasets at 3 months reported by MAJIQ. Splicing events having percent spliced-in (PSI) < 0.05 in *Fus*^*+/+*^ are highlighted in dark blue. (**H**) Volcano plot showing splicing alterations in *Fus*^*-/-*^ versus *Fus*^*+/+*^ datasets reported by MAJIQ. (**I**) Scatter plot showing the correlation between the magnitude of splicing changes (ΔPSI) in *Fus*^*Δ14/Δ14*^ versus *Fus*^*+/+*^ (x-axis) and *Fus*^*-/-*^ versus *Fus*^*+/+*^ (y-axis) datasets. Red: junctions with increased PSI in both conditions; blue: junctions with decreased PSI in both conditions; black: junctions with non-concordant changes. (**J**) GO enrichment analysis results for genes with splicing alterations in *Fus*^*Δ14/Δ14*^ versus *Fus*^*+/+*^ datasets at 3 months. (**K**) GO enrichment analysis results for genes with splicing alterations in *Fus*^*-/-*^ versus *Fus*^*+/+*^ datasets at 3 months. (**L**) Western blot showing FUS levels in spinal cord lysates from *Fus*^*+/+*^, *Fus*^*+/Δ14*^ and *Fus*^*Δ14/Δ14*^ mice. The N-terminal FUS antibody detects both wild-type (higher band) and Δ14FUS (lower band). Total protein levels are used to ensure equal loading. (**M**) Quantification of FUS bands normalised to total protein levels in (L) highlights a dose-dependent increase in FUS protein levels in *Fus*^*+/Δ14*^ and *Fus*^*Δ14/Δ14*^ mice. (**N**) Schematic and sashimi plot showing a decrease in *Taf15* intron 7 reads in spinal cord samples from *Fus*^*Δ14/Δ14*^ mice compared to *Fus*^*+/+*^ controls. (**O**) Schematic and sashimi plot showing a decrease in *Ewsr1* intron 8 and 9 reads in spinal cord samples from *Fus*^*Δ14/Δ14*^ mice compared to *Fus*^*+/+*^ controls.

**Fig. S2.**
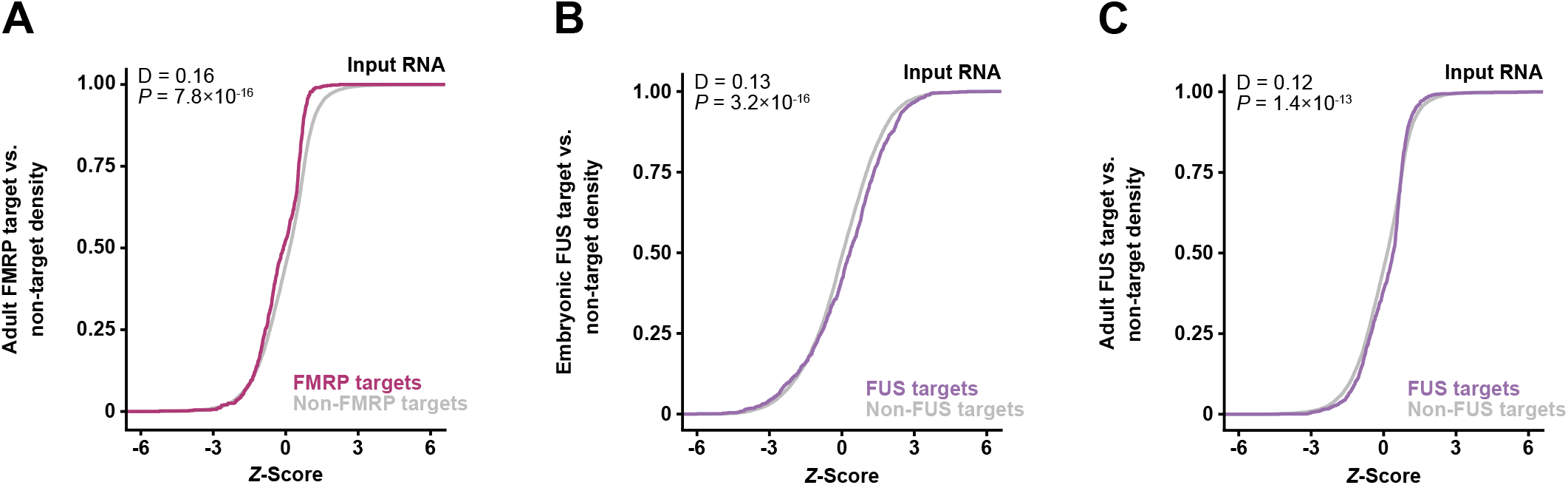
FMRP and FUS target expression analysis. (**A**) Cumulative frequency plot of *Z*-scores show a significant decrease in the expression of FMRP targets (plum) compared to non-target RNAs (grey) with similar expression levels in total *Fus*^*Δ14/Δ14*^ versus *Fus*^*+/+*^ adult spinal cord samples (*P* = 7.8 × 10^−16^, Kolmogorov-Smirnov). (**B**) Cumulative frequency plot of *Z*-scores show an increase in the expression of FUS targets (purple) compared to non-target RNAs (grey) with similar expression levels in embryonic (E17.5) total *Fus*^*Δ14/Δ14*^ versus *Fus*^*+/+*^ spinal cord samples (*P* = 3.2 × 10^−16^, Kolmogorov-Smirnov). (**C**) Cumulative frequency plot of *Z*-scores show a slight increase in expression of FUS targets (purple) compared to non-target RNAs (grey) with similar expression levels in total *Fus*^*Δ14/Δ14*^ versus *Fus*^*+/+*^ adult spinal cord samples (*P* = 1.4 × 10^−13^, Kolmogorov-Smirnov).

**Fig. S3.**
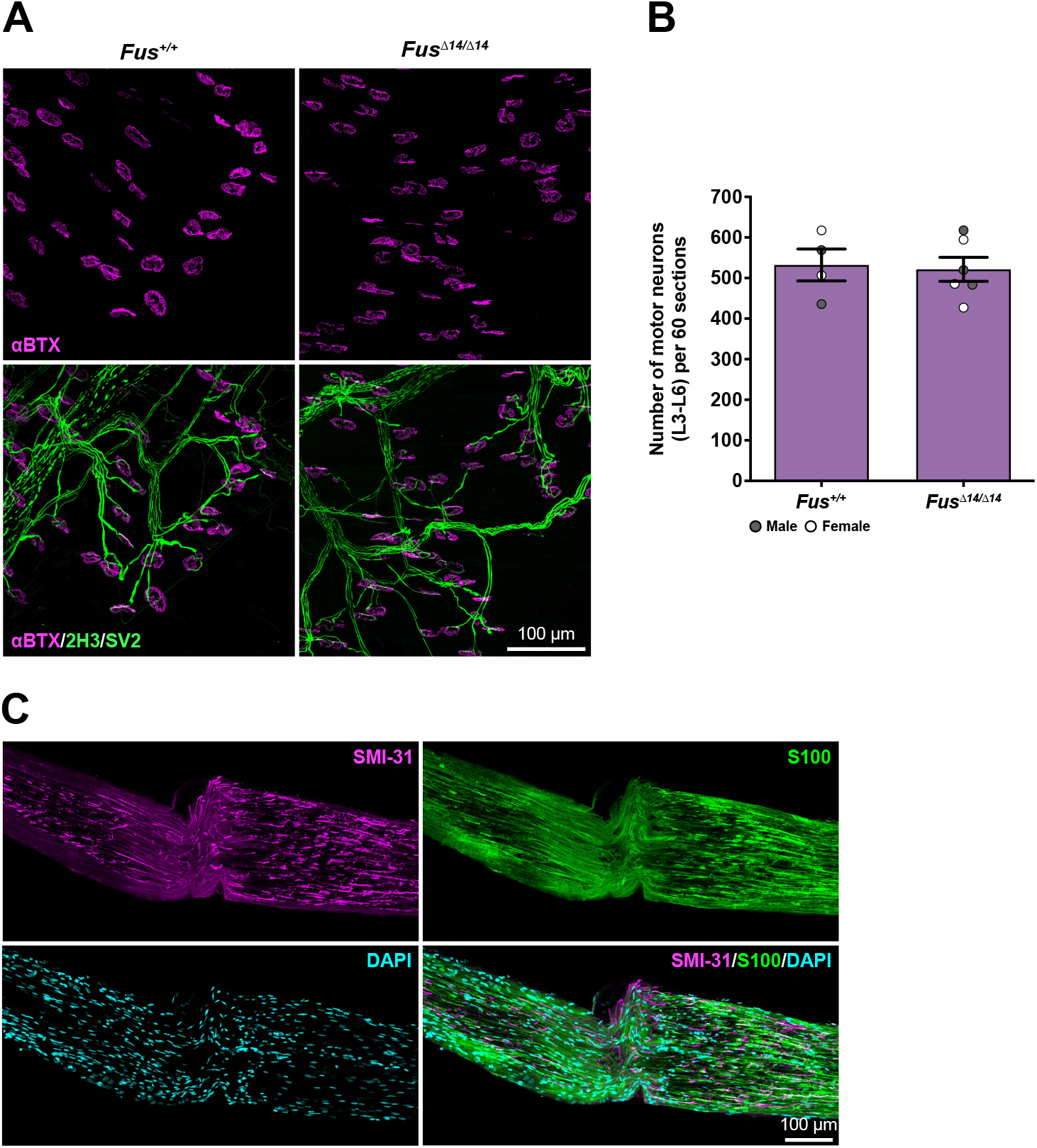
Δ14 FUS expression impacts endplate size, but does not affect motor neuron counts. (**A**) Representative maximum z-stack projection images of neuromuscular junctions from *Fus*^*+/+*^ (left) and *Fus*^*Δ14/Δ14*^ (right) mice. Postsynaptic acetylcholine receptor labelling (αBTX, magenta) was used to quantify endplate area. (**B**) *Fus*^*+/+*^ and *Fus*^*Δ14/Δ14*^ mice present a similar number of motor neurons in the lumbar spinal cord (*P* = 0.83, unpaired *t*-test). *n* = 4-6. (**C**) Representative maximum z-stack projection image of the damaged sciatic nerve immediately after crush. Longitudinal sections of 15 µm were stained with anti-S100 (green), anti-SMI-31 (magenta) and DAPI (blue) to visualise Schwann cells, phospho-neurofilament H and nuclei, respectively. Scale bars = 100 µm.

**Fig. S4.**
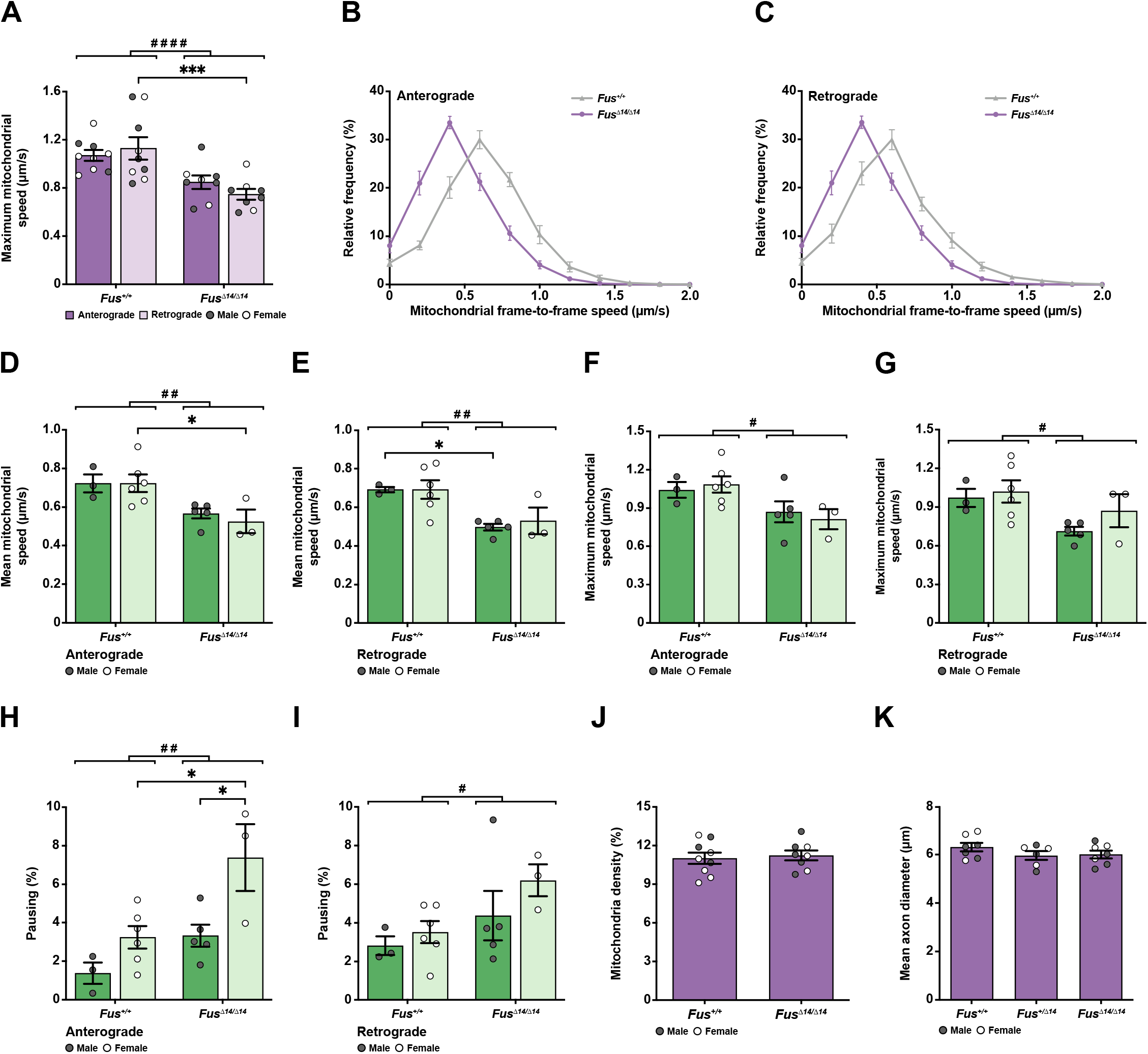
*In vivo* anterograde and retrograde axonal transport of mitochondria is impaired in *Fus*^*Δ14/Δ14*^ mice. (**A**) Mitochondrial transport analysis reveals decreased maximum speeds of anterogradely and retrogradely moving mitochondria in *Fus*^*Δ14/Δ14*^ sciatic nerve axons compared to *Fus*^*+/+*^ controls (*P* < 0.0001 for genotype, *P* = 0.75 for direction, *P* = 0.23 for interaction; two-way ANOVA). *n* = 8-9. (**B,C**) Frame-to-frame speed distribution curves of anterogradely (B) and retrogradely (C) moving mitochondria in *Fus*^*Δ14/Δ14*^ mice (purple) show a leftward shift compared to *Fus*^*+/+*^ (grey) animals, indicating a slower transport profile. (**D,E**) The decrease in mean mitochondrial transport speed for anterogradely (D) and retrogradely (E) moving organelles in *Fus*^*Δ14/Δ14*^ mice is similar between males and females (anterograde: *P* = 0.002 for genotype, *P* = 0.67 for sex, *P* = 0.66 for interaction; two-way ANOVA; retrograde: *P* = 0.002 for genotype, *P* = 0.71 for sex, *P* = 0.72 for interaction; two-way ANOVA). *n* = 3-6. (**F,G**) The decrease in maximum mitochondrial transport speed for anterogradely (F) and retrogradely (G) moving organelles in *Fus*^*Δ14/Δ14*^ mice is comparable between males and females (anterograde: *P* = 0.01 for genotype, *P* = 0.93 for sex, *P* = 0.53 for interaction; two-way ANOVA; retrograde: *P* = 0.03 for genotype, *P* = 0.25 for sex, *P* = 0.53 for interaction; two-way ANOVA). *n* = 3-6. (**H**) Anterogradely moving mitochondria spend a greater percentage of time paused in *Fus*^*Δ14/Δ14*^ mice compared to *Fus*^*+/+*^controls, with significantly more pausing occurring in females than males (*P* = 0.003 for genotype, *P* = 0.004 for sex, *P* = 0.22 for interaction, two-way ANOVA). *n* = 3-6. (**I**) Retrogradely moving mitochondria spend a greater percentage of time paused in *Fus*^*Δ14/Δ14*^ mice compared to *Fus*^*+/+*^ controls, with no significant difference between sexes (*P* = 0.05 for genotype, *P* = 0.22 for sex, *P* = 0.58 for interaction; two-way ANOVA). *n* = 3-6. (**J**) Mitochondria density is constant between *Fus*^*Δ14/Δ14*^ mice and *Fus*^*+/+*^ controls (*P* = 0.72, unpaired *t*-test). *n* = 8-9. (**K**) Axonal diameter, measured using the signal of HB9-driven GFP expression, is consistent between *Fus*^*+/+*^/*HB9::GFP, Fus*^*+/Δ14*^/*HB9::GFP* and *Fus*^*Δ14/Δ14*^/*HB9::GFP* mice (*P* = 0.35, Kruskal-Wallis). *n* = 6-7. For all graphs, \**P* < 0.05, \*\*\**P* < 0.001 with Šídák’s multiple comparisons. # shows comparison between genotypes. Grey circles = males, white circles = females.

